# Insulin receptor trafficking and interactions in muscle cells

**DOI:** 10.1101/2021.06.29.450241

**Authors:** Haoning Howard Cen, Aurora J. Mattison, Alireza Omidi, Jason Rogalski, Libin Abraham, Guang Gao, Michael R. Gold, Leonard J. Foster, Jörg Gsponer, James D. Johnson

## Abstract

Insulin resistance contributes to type 2 diabetes and can be driven by hyperinsulinemia. Insulin receptor (INSR) internalization and cell-surface dynamics at rest and during insulin exposure are incompletely understood in muscle cells. Using surface labelling and live-cell imaging, we observed robust basal internalization of INSR in C2C12 myoblasts, without an effect of added insulin. Mass-spectrometry using INSR knockout cells as controls, identified high-confidence binding partners, including proteins associated with internalization. We confirmed known interactors, including IGF1R, but also identified underappreciated INSR-binding factors such as ANXA2. AlphaFold-Multimer analysis of these INSR-binding proteins predicted potential INSR binding sites of these proteins. Protein-protein interaction network mapping suggested links between INSR and caveolin-mediated endocytosis. INSR interacted with both caveolin and clathrin heavy chain (CLTC) in mouse skeletal muscle and C2C12 myoblasts. Whole cell 2D super-resolution imaging revealed that high levels of insulin (20 nM) increased INSR colocalization with CAV1 but decreased its colocalization with CLTC. Single particle tracking confirmed the colocalization of cell-surface INSR with both over-expressed CAV1-mRFP and CLTC-mRFP. INSR tracks that colocalized with CAV1 exhibited longer radii and lifetimes, regardless of insulin exposure, compared to non-colocalized tracks, whereas insulin further increased the lifetime of INSR/CLTC colocalized tracks. Overall, these data suggest that muscle cells utilize both CAV1 and CLTC-dependent pathways for INSR dynamics and internalization.

## Introduction

The insulin receptor (INSR) is one of the most studied membrane proteins due to its importance in insulin resistance and diabetes. However, many fundamental aspects of INSR biology remain unclear, including the mechanisms and kinetics of its internalization. The INSR can be internalized via endocytosis (1,2), and endosomal INSR localization contributes to the phosphorylation of specific downstream targets when compared with INSR at the cell surface (3,4). Moreover, ligand-receptor complex dissociation in acidified endosomes permits insulin degradation, while the receptor can be sorted to recycling or degradation pathways (1,2). INSR internalization is reduced in monocytes(5,6) and adipocytes (7) obtained from patients with insulin resistance and type 2 diabetes, suggesting the importance of this process in disease pathogenesis.

Tissue-specific insulin resistance has been implicated in the development of diabetes (8,9), but it is less clear whether there are tissue-specific INSR internalization kinetics or mechanisms. Interestingly, adipocytes internalize insulin at a lower rate in the presence of high insulin concentrations, as compared to low insulin conditions (10), while hepatocytes are the opposite (11). The consensus view is that INSR is endocytosed via a clathrin-coated pit mechanism, based on evidence from a variety of cell types, including human and rat hepatocytes, fibroblasts, and lymphocytes (11–15). However, other studies have shown that caveolin-rich membrane lipid rafts, caveolae, are critical for INSR internalization in other cell types, including adipocytes (16,17). In pancreatic β-cells, we showed that caveolin-1 (CAV1) mediates INSR internalization, bypassing clathrin positive compartments (4). However, INSR internalization kinetics and routes are less well characterized in muscle cells, despite that muscle is one of the major insulin target tissues. It is estimated that skeletal muscle is a major glucose depot accounting for 60-70% of glucose uptake in response to insulin during hyperinsulinemic-euglycemic clamp(18). Under postprandial conditions, muscle disposes around 1/3 of orally derived glucose(19). In muscle cells, one study has shown that clathrin-mediated endocytosis is involved in INSR internalization(20), but the role of the caveolin-mediated pathway has not been investigated.

In the present study, we aim to characterize the dynamics of INSR internalization and endocytosis routes in muscle cells. We employed inter-domain tagged INSR with fluorescent proteins (4) and SNAP-tag (21), surface biotinylation assays, as well as super-resolution and TIRF imaging, to explore the dynamics of INSR internalization and the effects of ambient insulin. We also used proteomics and AlphaFold-Multimer to identify INSR interacting proteins and defined the relationship between INSR and caveolin or clathrin. Together, our data show that INSR exhibited a high degree of insulin-independent internalization in C2C12 myoblasts.

## Results

### Effects of insulin on INSR internalization

First, we characterized INSR internalization in C2C12 myoblasts under basal conditions and in the context of added insulin. Circulating insulin in humans oscillates in a range between approximately 0.01 nM and 0.75 nM (22–27). As for mice, fed insulin is ∼0.2 nM in lean mice and ∼3-5 nM in extremely obese mice (28). We used insulin concentrations of 0.2, 2, or 20 nM to cover a spectrum from physiological to supraphysiological concentrations. A higher rate of basal INSR internalization was observed in high-glucose serum-free DMEM medium when compared with PBS. The net INSR internalization over 15 min under various insulin concentrations in the DMEM medium did not significantly differ from the basal condition (Fig. 1A). Therefore, INSR has a high rate of constitutive internalization in this cell type, which may be enhanced by the nutrients in the medium, but that is largely independent of acute exposure to insulin across a range of physiological doses. These observations in myoblasts are consistent with our previous measurements in differentiated C2C12 myotubes, in which INSR internalization in insulin-free DMEM media is evident; insulin-stimulated internalization over time is also obvious but does not significantly differ at different insulin doses (29).

**Figure 1.**
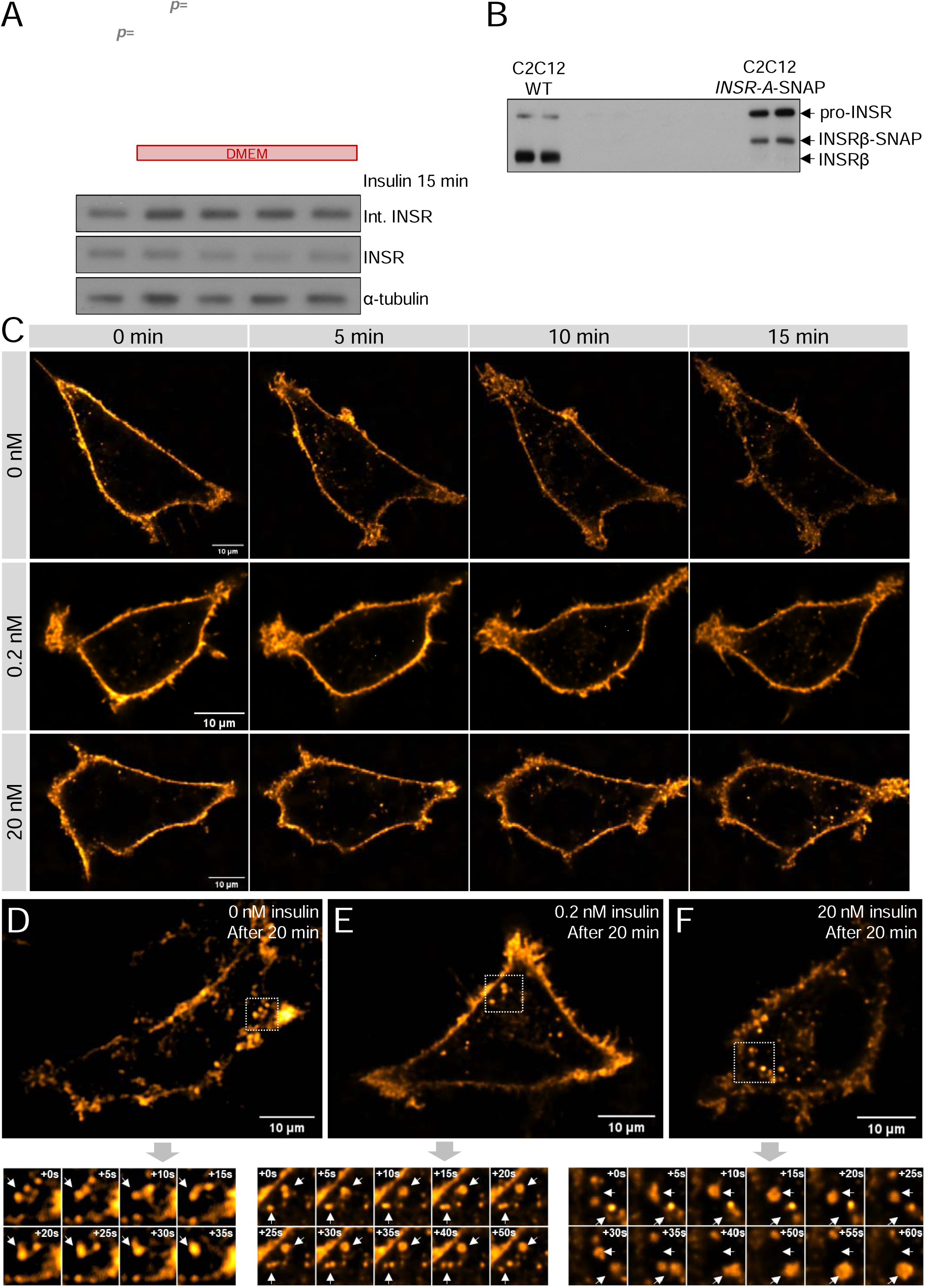
Biochemical and live-cell imaging characterization of the effects of insulin on INSR. **(A)** Internalized to total INSR ratio was quantified using surface biotinylation in undifferentiated C2C12 myoblasts. Cells were incubated in PBS (no insulin) or serum-free DMEM containing 0, 0.2, 2, or 20 nM insulin for 15 min. The ratios are normalized to the 0nM group of each gel. (p>0.05 when not specified.) **(B)** Western blot showing SNAP-tagged INSR expressed from lentiviral vector in comparison to wild-type INSR in C2C12 myoblasts. **(C,D)** Representative images of INSR-A-SNAP -labelled using an Alexa Fluor 488 cell non-permeable dye in 0, 0.2 or 20 nM insulin conditions in live undifferentiated C2C12 myoblasts from **(C)** 0 - 15 min or **(D)** after 20 min using a spinning disk confocal microscope. Time-lapse images of INSR vesicle interactions starting from the selected subregions (white squares) of snapshots are shown in the *insets*. (n=3 cells)

### Qualitative analysis of INSR kinetics in the context of different insulin doses

We also visualized INSR internalization at single-cell level using inter-domain-tagged INSR, which we have previously found more faithfully recapitulates the sub-cellular localization of INSR compared to terminal fusion proteins (4). The inter-domain-tagged INSR was also shown to have functional AKT and ERK signaling in C2C12 (Fig. S1 (30)) and other cell lines (4). C2C12 myoblast cells with stable overexpression of SNAP-tagged INSR isoform A (INSR-A-SNAP; Fig. 1B) were labelled using a cell non-permeable SNAP-Surface Alexa Flour 488 dye (Fig. 1C). Although a few INSR vesicles seemed to be internalized during the small gap between labelling and imaging, and were present at time zero, substantially more internalized INSR vesicles were observed at later timepoints (Fig. 1C). Live-cell time-lapse imaging of surface-labelled INSR-A-SNAP at the middle plane of cells by spinning disk confocal microscopy revealed INSR internalization in the context of 0, 0.2, and 20 nM insulin over 15 min (Fig. 1C, Video S1 (30)). Consistent with the surface biotinylation assay results in Figure 1A, we did not observe a clear difference in INSR internalization in the presence of insulin in these cells. Internalized INSR vesicles exerted dynamic movement and interaction, which was more evident between 20-30 min (Fig. 1D-F, Video S2 (30)). INSR-containing vesicles exhibited many apparent fusion and fission events regardless of insulin concentrations. Some smaller vesicles merged into a larger vesicle that remained in the field, undergoing limited and apparently random motion (one example is shown in Fig. 1F), suggestive of a fusion event. A caveat associated with 2D confocal imaging is limited Z resolution, raising the possibility that the observed INSR vesicles are not all in the same optical plane. Our experiments using surface labelling of SNAP-tagged INSR permitted the visualization of the dynamic behaviour of internalized INSR vesicles, and these qualitative observations motivated the more quantitative study below.

### Quantitative analysis of INSR kinetics in the context of insulin

Lateral movement of tyrosine kinase receptors has been linked to their functional state. Ligand binding of some monomeric receptors, such as EGFR, promotes their dimerization and aggregation and decreases their lateral mobility (31). Although INSRs are thought to already exist as dimers in plasma membrane, aggregation of INSR by antibodies can stimulate INSR signaling (31), and INSR may need to localize to membrane lipid rafts for normal signaling (32,33). Therefore, we sought to quantitatively assess whether insulin affects the lateral movement of INSR in or near the plasma membrane. To do this, we used time-lapse TIRF microscopy of C2C12 myoblast transiently expressing inter-domain-tagged INSR-A-EGFP. We imaged the same cells at 0, 6, or 11 min in the presence of 2 nM insulin, as well as a control group without insulin to control for other variables such as photobleaching (Fig. 2A). Single-particle tracking of INSR-A-EGFP (Fig. 2B) was used to calculate their diffusion coefficient (Fig. 2C,D) and track radius (Fig. 2E,F). Insulin did not have a big impact on the track radius or diffusion coefficient, despite some very subtle differences in the cumulative probability curves with very close median values (Fig. 2D,F). Together with the qualitative data reported above, these observations suggest that INSR mobility is largely unaffected by acute insulin stimulation in this cell model.

**Figure 2.**
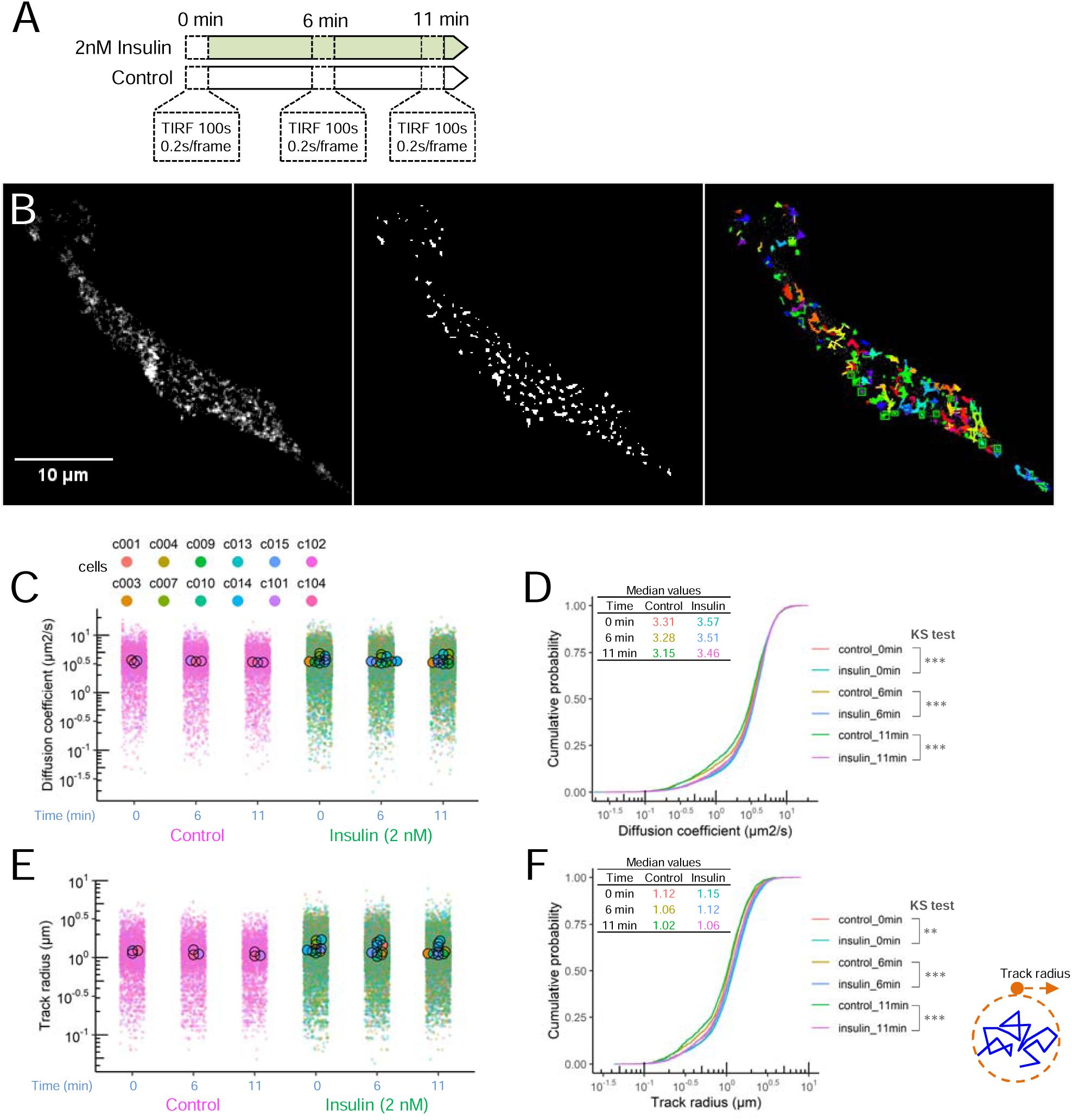
Live-cell TIRF imaging of cell-surface INSR. **(A)** Experimental design of TIRF microscopy of the same cells at 3 timepoints with 2 nM insulin or no insulin (control). **(B)** Representative TIRF image of INSR-A-EGFP puncta, binary image of detected INSR spots, and INSR tracks. **(C)** Diffusion coefficient of INSR-A-EGFP tracks in control or insulin group. Data is plotted as SuperPlot with larger dots showing mean values of the cells and smaller dots showing individual INSR tracks of the cells. **(D)** The cumulative probability shows the distribution of diffusion coefficients in **(C)**. **(E)** Track radius of INSR-A-EGFP tracks in control or insulin group. **(F)** The cumulative probability shows the distribution of diffusion coefficient in **(E)**. (n=3 cells in control group, n=9 cells in insulin group)

### Novel INSR interactors identified by immunoprecipitation and mass spectrometry

To explore the molecular mechanisms associated with INSR internalization, we conducted co-immunoprecipitation followed by mass-spectrometry to identify INSR interactors. We used INSR knockout C2C12 myoblasts as a negative control and an INSR antibody that does not cross-react with Insulin-like growth factor 1 receptor (IGF1R)(34) to identify INSR interactors with higher specificity than any previously reported study. After filtering the detected proteins against the INSR knockout cell control lysates, we identified INSR and five high-confidence INSR-binding proteins (Fig. 3A). The immunoprecipitated abundance of these detected interactors was not significantly changed by the addition of 2 nM insulin (Fig. 3A). INSR is well established to form heterodimers with IGF1R(35–37), and its immunoprecipitation represented a type of internal positive control for this study. The other detected proteins are not well-known INSR interactors. To discover whether any of the identified INSR-binding proteins might be connected with endocytosis machinery, protein-protein interaction (PPI) network was constructed between these co-immunoprecipated proteins and all the proteins under the “Endocytosis” pathway in Kyoto Encyclopedia of Genes and Genomes (KEGG). We manually extracted a subnetwork that connected all our identified proteins, except for HIST1H1B which did not connect with any proteins in this network (Fig. 3B). In the resultant PPI network, annexin A2 (ANXA2) is connected with CAV1 and CAV3 (Fig. 3B), and it has been reported to localize in caveolae (38). Ribosomal protein SA (RPSA) is a laminin receptor reported to localize in plasma membrane lipid rafts (39). Heat shock protein B1 (HSPB1; a.k.a. HSP27) is exclusively found in insulin-responsive tissues and has been implicated in insulin sensitivity and INSR/IGF1R signalling (40–42). The PPI network and cellular localization of these interactors pointed to the possibility that INSR may also associate with caveolins and/or related proteins in myoblasts.

**Figure 3.**
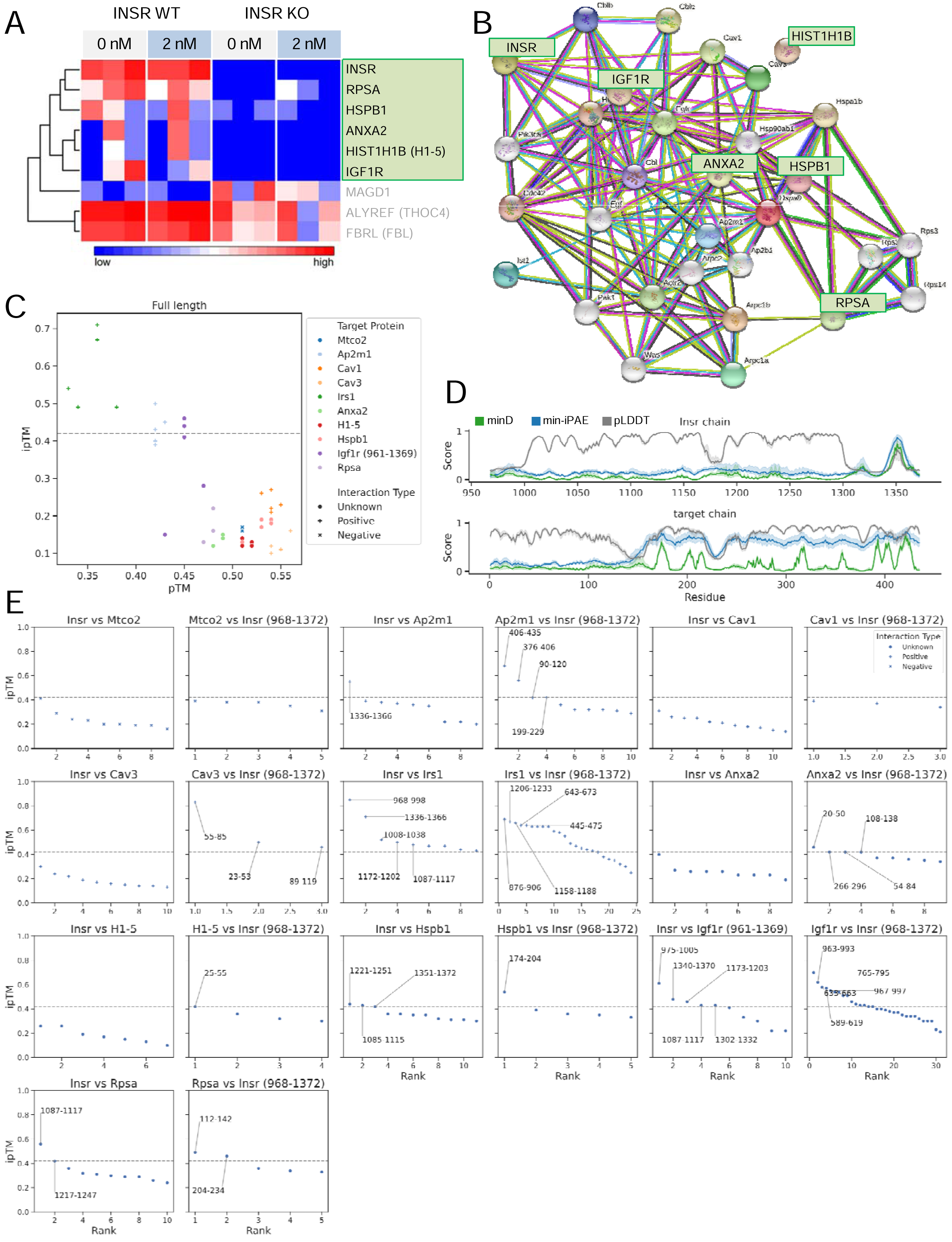
Identification of INSR interactors. **(A)** Proteins that are detected in INSR immunoprecipitation mass-spectrometry (IP-MS), and significantly different between wild-type and INSR knockout groups. Proteins in green area were specific interactors that had little detection in knockout cells. The other 3 proteins are also significantly different but not lower or absent in INSR KO cells. (n=3) (B) Protein-protein interaction network of INSR interactors and endocytosis proteins. **(C)** Modeling the interactions between the cytoplasmic region of INSR (residues 968 - 1372) and full-length interactor (target) proteins using AlphaFold-Multimer. pTM scores measure the accuracy of the overall structure of the protein complex and is relatively insensitive to localized inaccuracies. ipTM measures the accuracy of the interacting subunits of the complex, and ipTM > 0.42 (dash line) predicts direct interaction. The target proteins are classified as IP-MS identified novel interactors (Unknown ●), the known or expected interactors (positive **+**), and an expected non-interactor protein as a negative control (negative **×**). The cytoplasmic region of IGF1R (residues 961-1369) is used. (n = 5 predictions.) **(D)** Potential interaction sites based on per-residue scores along the INSR cytoplasmic region (residues 968 to 1372; top panel) and interactor (target) proteins Ap2m1 (bottom). minD (more accurate) and min-iPAE scores detect interaction sites. pLDDT shows whether the region is structured or unstructured. Shades show confidence errors of 99%. **(E)** ipTM scores for fragment-protein interactions. Thirty residues around the minD peaks (potential interacting sites) in **(D)** and **Fig. S2** (30) are analyzed. Each box relates to the interaction of the fragments of first protein vs the full-lenght second protein. For example, title “Insr vs Mtco2” means we have cut Insr into fragments and we have run AlphaFold-Multimer on each of those fragments vs full-length Mtco2. Each dot is the top ipTM score for one fragment (we do 5 predictions for each fragment vs protein). The protein residue positions of the interacting fragments (ipTM > 0.42) are labeled.

### Modelling INSR interactions *in silico* using AlphaFold-Multimer

AlphaFold-Multimer has been used to accurately model protein complexes (43,44). Recent work showed that AlphaFold-Multimer more accurately predicts interactions of protein fragments than lengthy full proteins and established a new pipeline to identify potential interacting fragments/sites (45). We applied this new pipeline to infer whether and how the putative INSR interactors identified with mass spectroscopy directly bind to INSR. We included a mitochondrial protein MTCO2 (aka COX2), expected to have no interaction with INSR, as a negative control in these *in silico* experiments. Several known or expected interactor of INSR were also tested, including IRS1, AP2M1, CAV1 and CAV3. First, we ran AlphaFold-Multimer on the cytoplasmic region of INSR and each of the full-length interactors to predict a protein complex structure. We made five separate predictions for each pair of proteins, shown as their predicted template modeling (pTM) and interface-pTM (ipTM) scores (Fig. 3C). Positive interaction with IRS1 and borderline interactions (ipTM ≥ 0.42 as suggested in (45)) with AP2M1 and IGF1R are predicted (Fig. 3C). CAV1, CAV3 and all other interactors did not have significant scores. The predictions with full-length proteins allowed us to extract per-residue scores of minimum expected distance (minD), minimum inter-predicted aligned error (min-iPAE), and the predicted local distance difference test (pLDDT) for INSR and all interactors (Fig. 3D and Fig. S2 (30)). minD and min-iPAE scores pinpoint potential interaction sites, with minD being the most accurate predictor of interaction sites between two proteins (45). pLDDT is a measure of local confidence and can indicate whether the region of interest is structured or unstructured. Using the INSR and AP2M1 interaction as an example, we observed that AlphaFold-Multimer predicts a sharp peak in all scores at residue 1350 of INSR, suggesting that the region around residue 1350 is in contact with AP2M1 (Fig. 3D). Yet this is not reflected in their full-length protein predictions (highest ipTM = 0.5, Fig. 3C), demonstrating that AlphaFold-Multimer may detect the interaction site between two proteins but is not able to predict the interaction when full-length proteins are used. To overcome this limitation, proteins can be fragmented into shorter segments, which are then submitted to AlphaFold-Multimer for prediction. This fragmentation can be guided by minD peaks that identify potential interaction sites. Thus, we cut both the INSR and the interactor proteins into fragments of 30 residues surrounding minD peaks (45). Doing so, we saw a large increase in ipTM scores for IGF1R, IRS1, AP2M1, CAV3, but not CAV1 (Fig. 3E). Interestingly, although we did not find positive predictions for CAV1 direct interaction, a peak in all scores (minD, min-iPAE and pLDDT) of CAV1 was detected around a phosphotyrosine residue 25, which might be an interacting site (Fig. S2 (30)).

Predictions via fragmentation also suggest interactions between INSR and ANXA2, H1-5, HSBP1, and RPSA (Fig. 3E). The ipTM scores of the negative control MTCO2 remained low, supporting the confidence in the positive predictions (Fig. 3E). Overall, the AlphaFold-Multimer predictions support the direct interactions between INSR and the novel interactors and identify potential interacting sites.

### Insulin-dependent interaction between INSR and CAV3 or CLTC *in vivo*

INSR has not been shown to utilize both caveolin and clathrin-mediated endocytosis in the same tissue; therefore, we sought to confirm that INSR could interact with both caveolin and clathrin in skeletal muscle *in vivo*. Skeletal muscle was collected 5 or 10 min after intraperitoneal injection of PBS or insulin (1.5 U/kg), an insulin dose and timepoints sufficient to measure acute insulin signalling via both AKT and ERK signalling pathways (Fig. 4A). Acute insulin administration did not change the amount of INSR that could be immunoprecipitated from skeletal muscle, nor did it alter the association between INSR and CLTC (Fig. 4B). However, 10 min after insulin injection, there was a significantly increased interaction between INSR and caveolin, specifically, the mature skeletal muscle-specific isoform CAV3 (46,47) (Fig. 4B). These data show that acute insulin signalling results in an increased association between INSR and CAV3 *in vivo*, but does not define the cellular context for this interaction.

**Figure 4.**
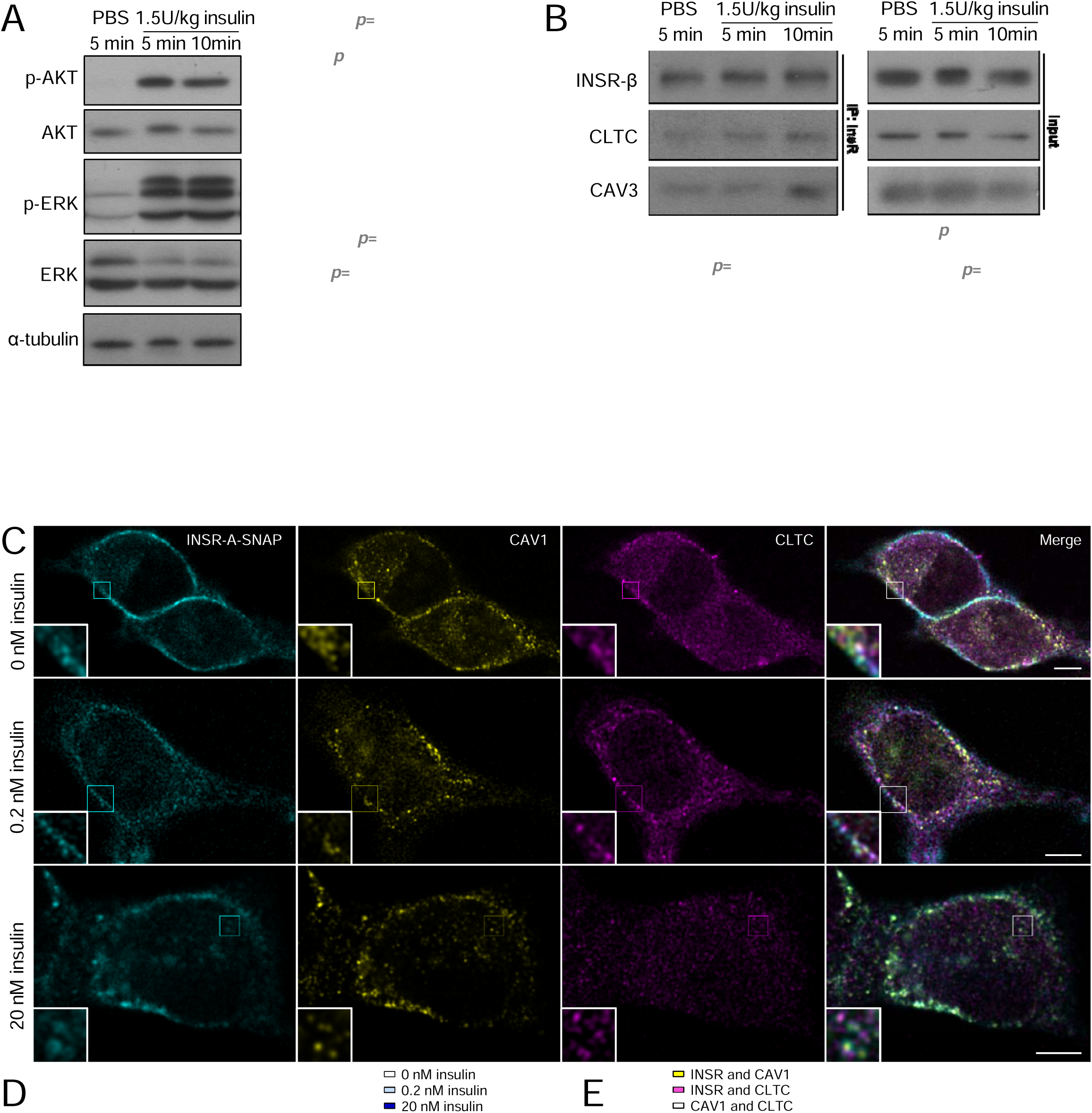
Interaction and colocalization between INSR, CAV3, CAV1 and CLTC under different insulin stimulations. **(A)** Western blot of mice skeletal muscle lysates after insulin injection. Phospho-AKT (Ser473) to total AKT ratio or phospho-ERK1/2 to total ERK ratio were quantified. (n=6) **(B)** Co-immunoprecipitation of skeletal muscle INSR after PBS or insulin injection. CLTC to INSR ratio or CAV3 to INSR ratio were quantified. (n=7-10) **(C)** Representative STED microscopy images of C2C12 myoblasts expressing INSR-A-SNAP (surface labeled) that were fixed after stimulation with 0 nM, 0.2 nM, or 20 nM insulin for 30 min and stained for CAV1 and CLTC (Scale bar = 5 µm). **(D,E)** Colocalizations between INSR-A-SNAP, CAV1, and CLTC were quantified by Object Pearson Coefficient. Data are plotted to show differences between insulin concentrations **(D)** or between protein pairs **(E)**. (n=7-9 images, 1-4 cells per image. *p<0.05, Tukey’s multiple comparison after 2-ANOVA. Box represents median and 25th to 75th percentiles.)

### Super-resolution imaging of INSR, CAV1, CLTC, and AXNA2 interactions in the presence of insulin

We next used super-resolution STED imaging to examine colocalization between INSR and clathrin (CLTC) or caveolin (C2C12 myoblasts only express CAV1 before differentiation (48)). C2C12 myoblast cells with stable overexpression of INSR-A-SNAP were labelled using a cell-nonpermeable SNAP substrate to image only surface-associated and newly internalized INSR. Cells were incubated with 0, 0.2 or 20 nM insulin for 30 min, and then fixed. Endogenous CLTC and CAV1 were then labelled using immunofluorescent antibody staining (Fig. 4C). The colocalization between INSR-SNAP, CAV1, and CLTC was quantified by Object Pearson Coefficient (Fig. 4D,E). The degree of colocalization between INSR and CAV1 was similar in the presence of 0 or 0.2 nM insulin but was higher under 20 nM insulin (Fig. 4D). However, INSR and CLTC colocalized more at 0.2 nM insulin, but less at 20 nM insulin, compared to 0 nM insulin (Fig. 4D). CAV1 and CLTC do not have known biological interactions; therefore, their low level of colocalization served as an internal negative control and, indeed, it did not change under different insulin levels (Fig. 4D). Using the same data, the relative colocalizations between INSR and CAV1 or CLTC were also considered at different insulin concentrations, further illustrating how high insulin concentrations drive INSR to colocalize with CAV1 (Fig. 4E). At 0.2 nM, insulin promoted INSR interaction with CLTC whereas 20 nM insulin redirected the INSR from CLTC to CAV1. Together, these data revealed the insulin-dependent interaction of INSR with both CAV1 and CLTC *in vivo* and *in vitro*.

Since our novel interactor ANXA2 is a target of insulin signaling (49,50), we also examined its insulin-dependent colocalization with INSR using the same SNAP labelling method and STED microscopy. Surface labeled INSR was allowed to internalize for 15 or 30 min in the presence of 0, 0.2, 2 or 20 nM insulin. Endogenous ANXA2 proteins were then labelled using antibody immunofluorescence. Colocalized INSR and ANXA2 puncta were observed in all conditions (Fig. 5A,C). The degree of colocalization quantified by Object Pearson Coefficient remained to be moderate in most conditions (lower than the INSR/CAV1 or INSR/CLTC interactions above) (Fig. 5B,D), except that 0.2 nM insulin seemed to decrease their colocalization at 30 min (Fig. 5D). This observation suggests that INSR and ANXA2 interactions are moderate and may be negatively regulated by insulin signalling.

**Figure 5.**
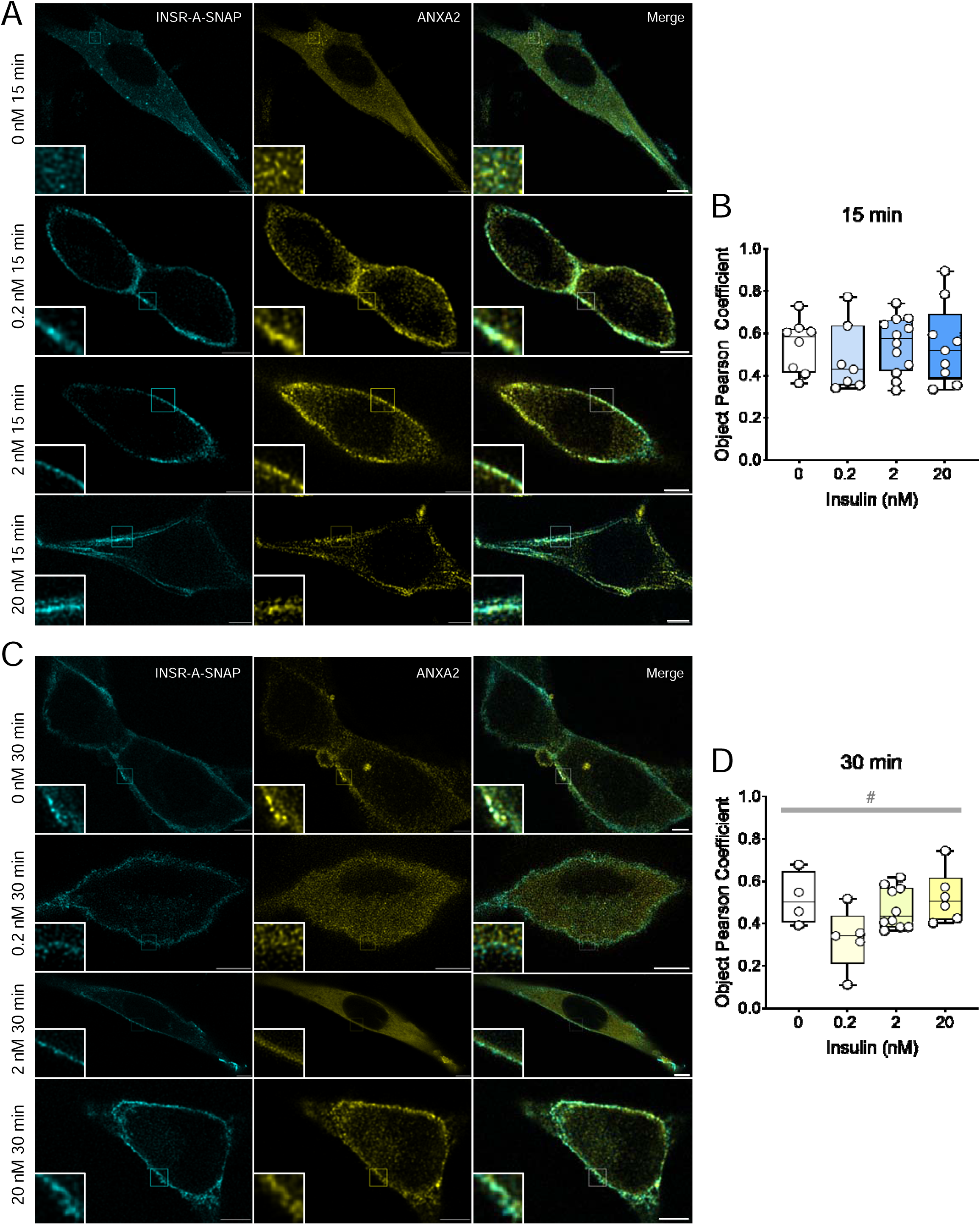
Colocalization and interaction between INSR and ANXA2 under different insulin concentrations. **(A,B)** Representative STED microscopy images of C2C12 myoblasts expressing INSR-A-SNAP (surface labeled) that were fixed after stimulation with 0 nM, 0.2 nM, or 20 nM insulin for (A)15 or (B)30 min and stained for ANXA2 (Scale bar = 5 µm). **(C,D)** Colocalization between INSR-A-SNAP and ANXA2 at (C)15 or (D)30 min of insulin-stimulated were quantified by Object Pearson Coefficient. (n=4-13 images, 1- 3 cells per image. ^#^p<0.05, 1-ANOVA of all groups. Box represents median and 25th to 75th percentiles.)

### Effects of CAV1 and CLTC on INSR mobility

To better understand the interplay between INSR, CAV1, and CLTC, we co-expressed INSR-B-TagBFP with CAV1-mRFP or CLTC-mRFP in C2C12 myoblasts and imaged them by TIRF time-lapse microscopy. Colocalizations between INSR and CAV1 or CLTC were observed (Fig. 6A,B). Some colocalized puncta moved or disappeared together with closely associated intensity, referred to here as colocalized tracks (examples in Fig. 6C,D). Tracks were considered as colocalized when the distance between the centers of particle signals was less than 2 pixels for more than 4 continuous frames. These observations suggested that INSR interacted with both CAV1 and CLTC on the cell surface.

**Figure 6.**
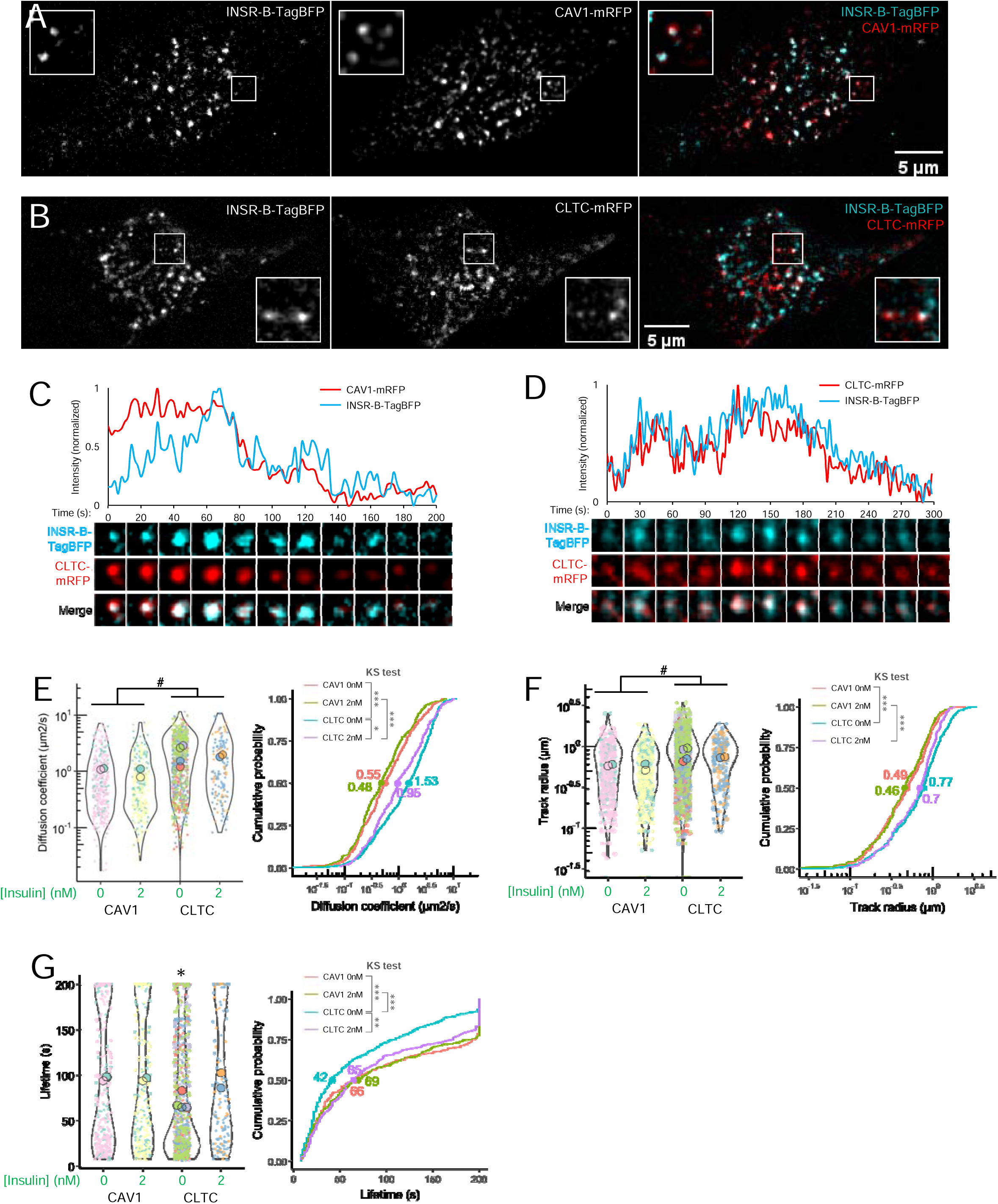
Live-cell TIRF imaging of cell-surface INSR, CAV1 and CLTC. **(A,B)** TIRF microscopy (2s/frame) showed colocalization between INSR-B-TagBFP and **(A)** CAV1-mRFP or **(B)** CLTC-mRFP. **(C,D)**The associated fluorescent intensity of representative INSR-B-TagBFP and colocalized **(C)** CAV1-mRFP or **(D)** CLTC-mRFP. **(E-G)** The (E) diffusion coefficient, (F) track radius, and (G) lifetime of INSR-B-TagBFP tracks in cells expressing CAV1-mRFP under 0 nM (n=2 cells, 360 tracks) or 2 nM insulin (n=2 cells, 324 tracks) or expressing CLTC-mRFP under 0 nM (n=4 cells, 917 tracks) or 2 nM insulin (n=2 cells, 146 tracks). Data is plotted as SuperPlot, with bigger dots showing mean values of the cells and smaller dots showing individual INSR tracks of the cells. The cumulative probability shows the distribution of the diffusion coefficient, with the dots and numbers showing the median values. (2-ANOVA: # effect of co-expression; post-hoc Tukey test: *p<0.05; Kolmogorov-Smirnov (KS) test: *p<0.05, **p<0.005, ***p<0.0005.) Experiments shown were conducted in Ringer’s buffer.

To define the effects of CAV1 or CLTC over-expression on INSR movement, we first conducted single particle tracking on all INSR puncta. Similar to our observations above (Fig. 2), insulin did not alter the diffusion coefficient or track radius of INSR. Cells with CAV1 co-expression had lower diffusion coefficients and track radius of INSR compared to cells with CLTC co-expression, regardless of ambient insulin exposure (differences in medians are at least ∼2 fold for diffusion coefficients and ∼1.5 fold for track radius; Fig. 6E,F). CLTC-expressing cells under 0 nM insulin had higher diffusion coefficients than those under 2 nM insulin (∼1.6 fold difference in median; Fig. 6E), as well as shorter INSR lifetimes than all other groups (>1.5 fold difference in median; Fig. 6G), which might suggest a more rapid internalization. Together, these data suggest that CAV1 may restrain INSR movement, relative to CLTC.

Finally, we performed an analysis within the same cells, comparing INSR tracks that were colocalized or not colocalized with either CAV1 or CLTC tracks, to further dissect the effects of CAV1 or CLTC. INSR appeared to spend more time associated with CAV1, and for longer periods, compared with CLTC (Fig. 7A,B). There were no clear effects of time on the association between INSR and CAV1 or CLTC across different moments of their lifetimes together (Fig. 7A,B). We compared the colocalized and the non-colocalized INSR tracks, for both CAV1 and CLTC, and found no significant differences in mean diffusion coefficients (Fig. 7C,D), except that in CLTC-expressing cells without insulin, non-colocalized INSR had higher diffusion coefficients than CLTC-colocalized INSR, based on the cumulative probability curve (Fig.7D). The track radius of INSR colocalized with CAV1 was larger than non-colocalized tracks regardless of insulin (Fig. 7E), but colocalization with CLTC had significant effects (Fig. 7F). Strikingly, the lifetimes of INSR tracks colocalized with CAV1 were much longer than non-colocalized tracks regardless of insulin (3.8∼4 fold differences in medians; Fig. 7G). INSR tracks colocalized with CLTC also had longer lifetimes than non-colocalized tracks (∼2.5 fold difference in median), and insulin also further increased the lifetime of colocalized tracks (Fig. 7H). Overall, our live-cell imaging of INSR confirmed its distinct interactions with either CAV1 or CLTC around the cell surface.

**Figure 7.**
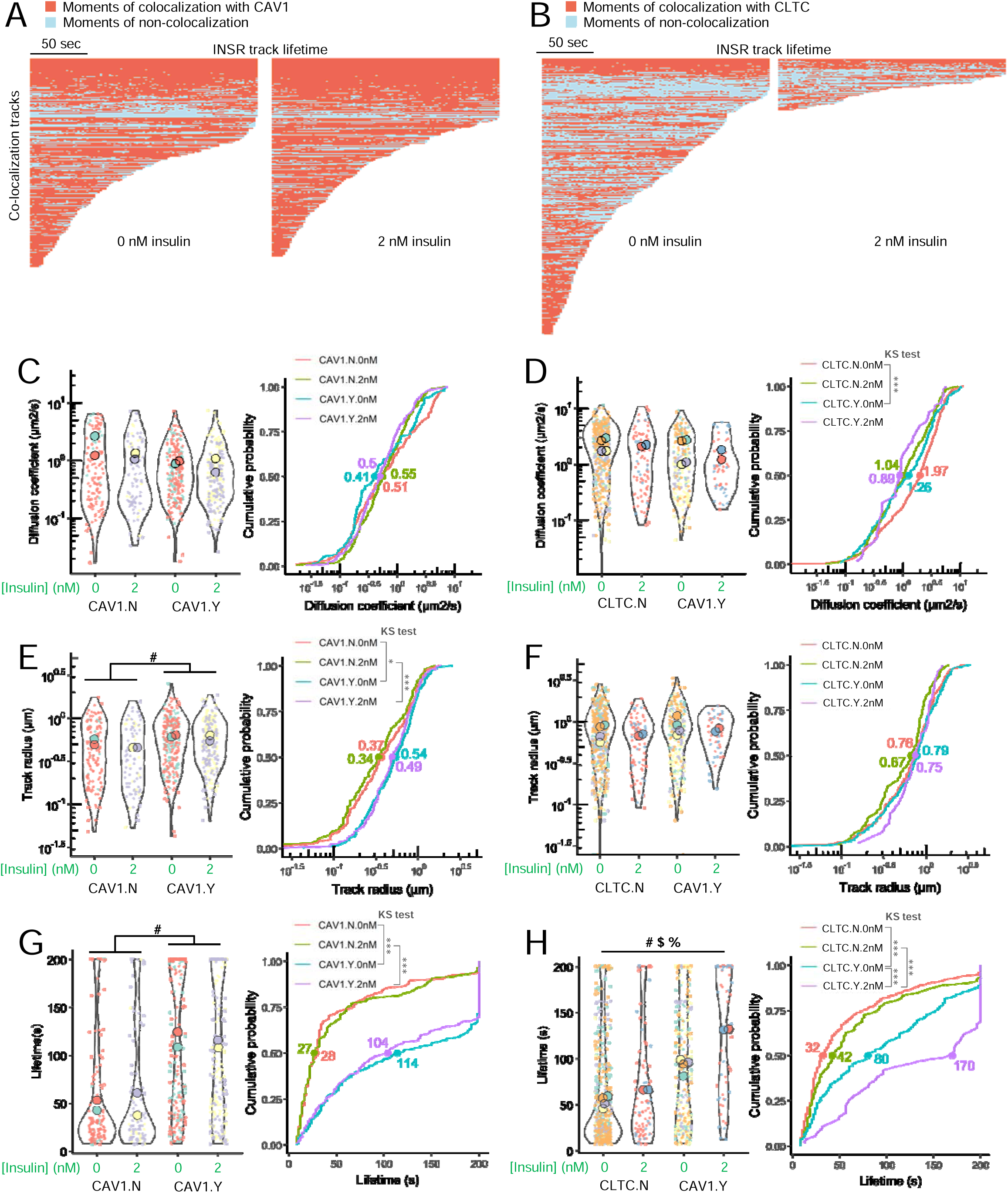
Analysis of INSR tracks colocalized or not colocalized with CAV1 or CLTC. **(A)** INSR tracks that colocalized with CAV1 under 0 or 2 nM insulin (referred to as CAV1.Y.0nM or CAV1.Y.2nM below). **(B)** INSR tracks that colocalized with CLTC under 0 or 2 nM insulin (referred to as CLTC.Y.0nM or CLTC.Y.2nM below). **(C)** Diffusion coefficient of INSR-B-TagBFP tracks that did not colocalize with CAV1-mRFP tracks under 0 nM insulin (CAV1.N.0nM) or 2 nM insulin (CAV1.N.2nM) or colocalized with CAV1-mRFP tracks under 0 nM insulin (CAV1.Y.0nM) or 2 nM insulin (CAV1.Y.2nM). The cumulative probability shows the distribution of the measurements, with the dots and numbers showing the median values (same for other cumulative probability plots). **(D)** Diffusion coefficient of INSR-B-TagBFP tracks that did not colocalize with CLTC-mRFP tracks under 0 nM insulin (CLTC.N.0nM) or 2 nM insulin (CLTC.N.2nM) or colocalized with CLTC-mRFP tracks under 0 nM insulin (CLTC.Y.0nM) or 2 nM insulin (CLTC.Y.2nM). **(E)** Track radius of INSR-B-TagBFP tracks that did not colocalize with CAV1-mRFP tracks under 0 nM insulin (CAV1.N.0nM) or 2 nM insulin (CAV1.N.2nM) or colocalized with CAV1-mRFP tracks under 0 nM insulin (CAV1.Y.0nM) or 2 nM insulin (CAV1.Y.2nM). **(F)** Track radius of INSR-B-TagBFP tracks that did not colocalize with CLTC-mRFP tracks under 0 nM insulin (CLTC.N.0nM) or 2 nM insulin (CLTC.N.2nM) or colocalized with CLTC-mRFP tracks under 0 nM insulin (CLTC.Y.0nM) or 2 nM insulin (CLTC.Y.2nM). **(G)** Lifetime of INSR-B-TagBFP tracks that did not colocalize with CAV1-mRFP tracks under 0 nM insulin (CAV1.N.0nM) or 2 nM insulin (CAV1.N.2nM) or colocalized with CAV1-mRFP tracks under 0 nM insulin (CAV1.Y.0nM) or 2 nM insulin (CAV1.Y.2nM). **(H)** Lifetime of INSR-B-TagBFP tracks that did not colocalize with CLTC-mRFP tracks under 0 nM insulin (CLTC.N.0nM) or 2 nM insulin (CLTC.N.2nM) or colocalized with CLTC-mRFP tracks under 0 nM insulin (CLTC.Y.0nM) or 2 nM insulin (CLTC.Y.2nM). (CAV1.N.0nM, n=2 cells, 125 tracks; CAV1.N.2nM, n=2 cells, 96 tracks; CAV1.Y.0nM, n=2 cells, 235 tracks; CAV1.Y.2nM, n=2 cells, 228 tracks; CLTC.N.0nM, n= 4 cells, 597 tracks; CLTC.N.2nM, n= 2 cells, 91 tracks; CLTC.Y.0nM, n= 4 cells, 320 tracks; CLTC.Y.2nM, n= 2 cells, 55 tracks.) (2-ANOVA: # effect of colocalization, $ effect of insulin, % interaction of the 2 variables; Kolmogorov-Smirnov (KS) test: *p<0.05, ***p<0.0005.) Experiments shown were conducted in Ringer’s buffer.

## Discussion

The goal of this study was to explore the INSR trafficking dynamics and interactions in a muscle cell model. We found that INSR was constitutively internalized both in the absence and presence of insulin. We used high-stringency proteomics and AlphaFold-Multimer to identify specific INSR binding proteins in this cell type and PPI network analysis to predict a role for caveolin in INSR internalization. Biochemical experiments, super-resolution imaging, and TIRF imaging suggested both caveolin and clathrin are involved in INSR internalization and cell-surface trafficking (Fig. 8).

**Figure 8.**
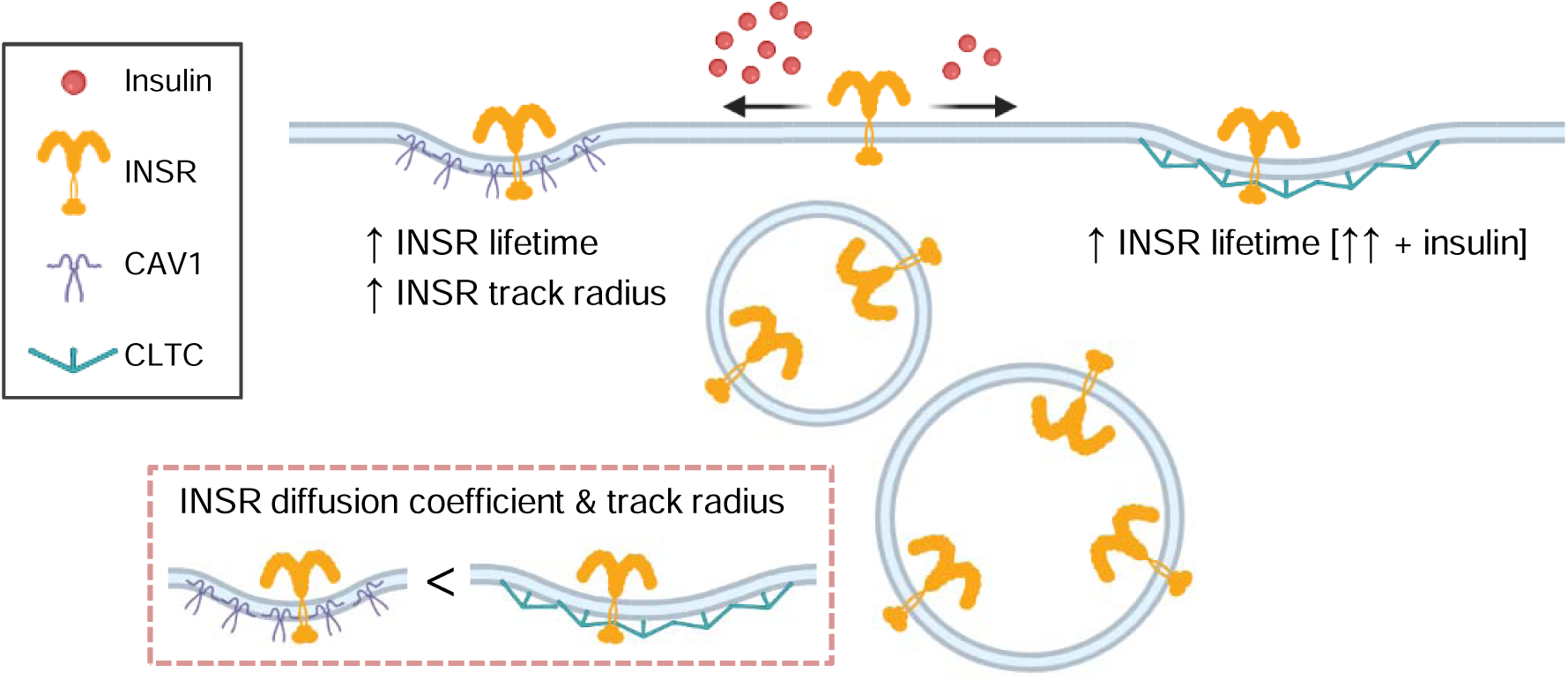
Summary of the effects of CAV1 and CLTC on INSR tracks. Higher insulin (20 nM) promoted INSR and CAV1 colocalization, while lower insulin (0.2 nM) promoted INSR and CLTC colocalization. INSR tracks had lower diffusion coefficient and track radius in CAV1-overexpressing cells than CLTC-overexpressing cells. Within the same cells, INSR tracks colocalized with CAV1 had longer lifetimes and larger track radius than non-colocalized tracks. INSR tracks colocalized with CLTC had longer lifetimes than non-colocalized tracks, which is further increased by insulin.

The constitutive internalization of INSR has been described in lymphocytes and shown not to require ligand binding or kinase activity (although insulin-stimulated autophosphorylation of INSR could accelerate this process (14,51)). In differentiated C2C12 myotubes, INSR phosphorylation levels were constitutively high, with little additional activation by insulin across the physiological range (29), consistent with the lack of stimulation of INSR internalization by acute insulin treatment in this study. Surface labeling by SNAP-tag enabled clear visualization of internalized INSR-containing vesicles, permitting the study of INSR endosomal sorting and traffic. The lateral movement of INSR in or near the plasma membrane was also not dramatically affected by insulin, consistent with a previous study that reported unchanged INSR diffusion coefficient with or without insulin in HEK293T, HepG2, and human IR-overexpressing Rat-1 cells (52). CAV1 may have a role in decreasing the mobility of INSR, perhaps due to their restricted lateral movement in the plasma membrane owing to actin cytoskeleton connections through Filamin-A (53). Accumulation of ganglioside GM3 in TNFα-induced insulin resistance was associated with the elimination of INSR from the caveolae and increased INSR mobility on the cell surface (32).

The possibility of more than one internalization pathway for INSR in some cell types has been understudied. Dual clathrin and caveolin endocytosis routes have been suggested for IGFR in Ewing sarcoma cells (54), which has a similar structure and function as INSR and can form heterodimers with INSR (55). The current study supports the concept that INSR may also utilize both clathrin- and caveolin-mediated endocytosis in the same cell type. In hepatocytes, it has been suggested that INSRs are internalized in two functionally separate pathways based on the much more rapid recycling of receptors internalized by high concentrations of insulin (17.2 nM) versus low insulin (0.17 nM); treatments known to inhibit coated pit formation blocked internalization of the insulin-INSR complex at low but not high insulin concentrations (11). Most studies of INSR internalization examined only one route. The mechanisms of coordinated multiple internalization routes, implied by the insulin-dependent clathrin and caveolin interactions we observed in muscle cells, might direct INSR to different destinations (e.g. recycle or degradation)(56) and should be comprehensively characterized in more cell types to assess potential roles in tissue-specific insulin resistance. Generally, we expect our results will translate to other muscle cell models and human muscle, given the strong evolutionary conservation of insulin signaling (57), but our results should be validated in human muscle in future studies.

We used stringent proteomics and INSR CRISPR knockout control cells to identify known, novel, and under-appreciated interactors of INSR. As expected, we identified IGF1R, an established hetero-dimerization partner of INSR. We also demonstrated the specific binding of HSBP1 (HSP27), which is involved in tyrosine kinase signalling pathways downstream of p38 MAPK, ERK, and MAPKAPK5 (58) and has been previously linked to insulin sensitivity (40–42). We demonstrated an interaction between INSR and RPSA, which has been previously co-immunoprecipitated with INSR and insulin receptor substrates in 3T3L1 preadipocytes (59). In other cell types, RPSA drives ERK signalling and cytoskeletal organization, with nuclear functions affecting cell growth, survival, migration, and protein synthesis (39). ANXA2 is not a known interactor of muscle INSR, but it has been reported in NIH3T3 fibroblasts (60). Other studies found that INSR signalling can tyrosine-phosphorylate (49,50) and SUMOylate (61) AXNA2, which mediates insulin-stimulated actin rearrangement. Interestingly, internalized INSR seemed necessary for AXNA2 phosphorylation because mutant INSR that was incapable of internalization but normal for tyrosine kinase signalling failed to phosphorylate AXNA2 (49). AXNA2 also interacts with endosomal and adaptor prortein APPL2, which interacts with INSR (62,63). The PPI network showed that ANXA2 has direct connection with the epidermal growth factor receptor (EGFR) which belongs to the same receptor tyrosine kinase superfamily as INSR (64). ANXA2 knockdown halted EGFR endocytosis beyond the early endosome, which was accompanied by enhanced EGFR and AKT phosphorylation (65). These data support the possibility that ANXA2 may have a role in INSR trafficking and signalling that is worth investigating. Interestingly, evidence compiled on the Common Metabolic Disease Knowledge Portal shows that ANXA2 genetic variation is strongly associated with fasting C peptide levels, pointing to a causal role, direct or indirect, in hyperinsulinemia/insulin resistance. Both *Anxa2* and *Rpsa* mRNAs were upregulated by prolonged insulin in C2C12 myotubes (8). HIST1H1B belongs to the family of linker histones that is important in chromatin organization (66), but the significance of this INSR interaction requires further investigation. Our stringent proteomic analysis, together with AlphaFold-Multimer modelling, provides evidence for a limited number of proteins that are likely to be direct interactors with INSR in myoblast cells, and demonstrates that the majority of these interactors were not significantly altered by insulin exposure.

There are limitations and unexplained observations in the current study. Although our inter-domain-tagged INSR has functional signalling and internalization, it tends to have higher pro-INSR bands (Fig. 1B)(4), which is indicative of a potential processing issue. Nonetheless, higher pro-INSR could also occur in wildtype C2C12 myoblasts (Fig. S1A (30)), possibly due to stress (67), which may be a co-culprit in the processing issue of the overexpressed tagged INSR. STED superresolution microscopy usually has a resolution of 40-50 nm and might be able to reach 20 nm resolution in the best scenario, and primary and secondary antibody labelling might also add ∼30nm distance and reduce the accuracy (68). TIRF microscopy has diffraction-limited lateral resolution (∼200 – 300 nm) with the thickness of the excitation depth around 100 nm (69). Therefore, the colocalization analyses may not confirm the interactions of the proteins. Nonetheless, the insulin-dependent colocalization observed in STED microscopy and the track colocalization over multiple frames observed in TIRF microscopy are evidence that the proteins had interactions instead of random colocalization. This is further supported by the AlphaFold-Multimer predictions. The CAV1- or CLTC-colocalized INSR tracks had longer lifetimes than non-colocalized tracks, which conflicted with the assumption that CAV1 or CLTC internalized INSR, but instead, they seemed to retain INSR on the membrane. There are multiple possible explanations; for example, overexpressed CAV1 might stabilize caveolae (70), or CLTC could form stable flat lattices, termed clathrin plaques (71), which might have novel involvement in INSR internalization or signalling.

As we did not perform loss-of-function studies to confirm the roles of caveolin and clathrin in INSR endocytosis, further research is needed to elucidate the roles of specific INSR internalization routes on insulin actions. For example, inhibition of clathrin-mediated endocytosis of INSR by knocking down LMBD1 in myoblasts increased activation of the INSR and AKT (20). Our lab discovered that reducing CAV1-dependent INSR internalization blocked the activation of ERK, but not AKT, signalling in pancreatic beta-cells (4). Inhibition of INSR internalization in adipocytes (72–74) or hepatocytes selectively blunted different signaling branches (12,75). More recent advances in INSR endocytosis mechanisms and consequences are reviewed in (76,77). For example, INSR endocytosis is found to be regulated by the spindle checkpoint proteins (MAD2, CDC20, BUBR1 and p31^comet^)(78,79) and SHP1-MAPK pathway (80) in the liver and by the inceptor protein in beta cells (81,82); decreased insulin-stimulated endocytosis or increased insulin-independent endocytosis by manipulating these proteins can promote or impair insulin signaling, respectively. Therefore, INSR internalization enables selective activation of downstream signalling pathways, and in a tissue-specific manner. Future study could be done on human muscle cell lines or biopsies to confirm the interactions and their functional roles.

In summary, this study provides important information on the internalization of INSR in myoblast cells and defines the stringent INSR interactome. We also provide evidence using a variety of methods that INSR interacts with both clathrin and caveolins in this cell model. The current study can serve as a stepping stone for more detailed research on how multiple internalization routes can provide cell-type customized regulations on INSR internalization.

## Materials and Methods Cell culture

The C2C12 mouse myoblast (ATCC cell line provided by Dr. Brain Rodrigues, University of British Columbia, Vancouver, Canada) was maintained in Dulbecco’s modified Eagle’s medium (DMEM, Gibco) supplemented with 10% (v/v) fetal bovine serum (FBS, Gibco), and 1% penicillin-streptomycin (100 μg/ml; Gibco). All downstream experiments were conducted after 6-hour serum starvation in serum-free medium (DMEM supplemented with 1% penicillin-streptomycin), except as otherwise indicated.

### Western blot analyses

C2C12 myotubes or mice skeletal muscle (gastrocnemius) tissues were sonicated in RIPA buffer (50 mM β-glycerol phosphate, 10 mM HEPES, 1% Triton X-100, 70 mM NaCl, 2 mM EGTA, 1 mM Na_3_VO_4_, and 1 mM NaF) supplemented with complete mini protease inhibitor cocktail (Roche, Laval, QC). Lysates were incubated in Blue Loading Buffer containing 50 mM DTT (Cell Signaling, Danvers, MA) at 95°C for 5 min and resolved by SDS-PAGE. Proteins were then transferred to PVDF membranes (BioRad, CA) and probed with antibodies against p-ERK1/2 (Thr202/Tyr204) (1:1000, Cat. #4370, RRID: AB_2315112), ERK1/2 (1:1000, Cat. #4695, RRID: AB_390779), p-AKT (Ser473) (1:1000, Cat. #9271, RRID: AB_329825), p-AKT (Thr308) (1:1000, Cat. #9275, RRID: AB_329828), AKT (1:1000, Cat. #9272, RRID: AB_329827), INSR-β subunit (1:1000, Cat. #3020S, RRID: AB_2249166), p-INSRβ (Tyr1150/1151) (1:1000, Cat. #3024, RRID: AB_331253), FOXO1 (1:1000, Cat. #2880, RRID: AB_2106495), p-FOXO1 (Thr24) (1:1000, Cat. #9464, RRID: AB_329842), all from Cell Signalling (CST), and β-tubulin (1:2000, Cat. #T0198, Sigma, RRID: AB_477556). The signals were detected by secondary HRP-conjugated antibodies (anti-mouse, Cat. #7076, RRID: AB_330924; anti-rabbit, Cat. #7074, RRID: AB_2099233; Cell Signaling) and Pierce ECL Western Blotting Substrate (Thermo Fisher Scientific). Protein band intensities were quantified with Image Studio Lite software (LI-COR).

### Surface Protein Biotinylation Assay

Biotinylation of surface proteins was performed as previously described(29). In brief, cells were incubated with cell-impermeable EZ-Link-NHS-SS-biotin (300 μg/ml in PBS; Pierce) at 37°C for 2 min and washed with ice-cold 50 mM Tris-buffered saline (TBS) to remove excess biotin. For detecting internalized proteins, cells were washed with PBS and incubated in PBS or serum-free medium supplemented with 0, 0.2, 2, or 20 nM insulin at 37°C to stimulate INSR internalization. After certain time periods, cells were incubated with glutathione solution (50 mM glutathione, 75 mM NaCl, 1 mM EDTA, 1% BSA, 75 mM NaOH) to strip the remaining surface biotin lysed in complete RIPA buffer. Biotinylated internalized proteins in lysates were captured by NeutrAvidin beads (Pierce) and eluted by incubating in Blue Loading Buffer (50 mM DTT, Cell Signaling, Danvers, MA) containing 50 mM DTT at 95°C for 5 min. Internalized INSR in eluent and total INSR in lysates were detected in Western blot analysis.

### Plasmids, SNAP-tag cloning and lentivirus infection

CAV1-mRFP and CLTC-mRFP plasmids were generous gifts from Dr. Ivan Robert Nabi, Vancouver, BC, Canada. INSR-A-TagBFP and INSR-B-TagBFP plasmids are previously described(4).

PCR amplification of DNA fragments was performed using Q5 High-Fidelity DNA Polymerase (Cat. #M0491, New England Biolabs, Ipswich, MA, USA) according to the manufacturer’s instructions. Prior ligation with the target vector, all PCR products were subcloned into pJET1.2 vector using the CloneJET PCR Cloning Kit (Cat. #K1232, Thermo Scientific, Waltham, MA, USA) according to the manufacturer’s instructions. Subsequently, the PCR products were excised from the pJET1.2 vector, gel purified and ligated with the digested target vector using T4 DNA ligase. To generate INSR-A-EGFP and INSR-B-EGFP plasmids, eGFP was PCR amplified using primers 5’-CGT ACG ATC GAA TTC CCC GTG AGC AAG GGC GAG GAG CTG-3’ (forward) and 5’- ACC GGT ATC TGA TCC GGA CTT GTA CAG CTC GTC CAT GCC GA-3’ (reverse) and inserted into pcDNA3.1(-)-INSR-A-TagRFP and pcDNA3.1(-)-INSR-B-TagRFP (previously described(4)) to replace TagRFP by EGFP using BsiWI and AgeI restriction sites.

To generate pLenti-INSR-A-SNAPf and pLenti-INSR-B-SNAPf lentiviral plasmids, SNAP tag was firstly PCR amplified from pSNAPf Vector (Cat. #N9183S, New England Biolabs, Ipswich, MA, USA) using primers 5’-CGT ACG ATC GAA TTC CCC ATG GAC AAA GAC TGC GAA ATG AAG CGC ACC ACC-3’ (forward) and 5’- ACC GGT ATC TGA TCC GGA ACC CAG CCC AGG CTT GCC CAG-3’ (reverse) and inserted into pcDNA3.1(-)-INSR-A-TagRFP and pcDNA3.1(-)-INSR-B-TagRFP to replace TagRFP by SNAP-tag using BsiWI and AgeI restriction sites. Subsequently, INSR-A-SNAP and INSR-B-SNAP ORFs were PCR amplified from pcDNA3.1(-)-INSR-A/B-SNAPf plasmids using primers 5’-GAA CCC ACT GCT TAC TGG CT-3’ (forward) and 5’-GGT GGT TGT ACA CTA GAA GGC ACA GTC GAG GC -3’ (reverse) and inserted into pLenti-C-Myc-DDK-IRES-GFP vector to replace the Myc tag, Flag tag, IRES, and TurboGFP using NheI and BsrGI restriction sites. The PCR program for this long INSR-SNAPf fragments was optimized to be as follows: initial denaturation 98°C 3min; 30 cycles of 98°C 10 s, 71°C 30 s, 72°C 4min; final extension 72°C 2min; hold 4°C.

### SNAP-tag labelling and spinning disk confocal imaging of C2C12 myoblast

Cell non-permeable SNAP-Surface Alexa Fluor 488 substrate (Cat. #S9129S, New England Biolabs) was dissolved and diluted to 5 µM concentration in serum-free medium containing 0.5% BSA according to the manufacturer’s instruction. Next, stable C2C12 myoblasts cell line expressing INSR-A-SNAP seeded on glass-bottom dishes (Cat. #P35G-1.5-14-C, MatTek) were incubated in the substrate for 30 min on ice and washed 3 times with cold FuoroBrite DMEM (Cat. #A1896701, Gibco) serum-free medium.

C2C12 dish was then set up on the Spinning disk microscope, applied 37°C FuoroBrite DMEM serum-free medium containing 0, 0.2 or 20 nM human insulin (Cat. #I9278, Sigma-Aldrich), and immediately started imaging in a 37°C chamber using a 100×/1.45 oil objective and QuantEM 512SC Photometrics camera on a spinning disk confocal system based on Zeiss Axiovert 200M microscope (LSI Imaging Core, UBC). A focal plane in the middle of the cell was acquired to avoid capturing the top and bottom of the cell. Time-lapse imaging was conducted at 30 s intervals between 0 – 20 min or at 5 s intervals between 20-30 min.

### Immunofluorescence staining

Stablly-transfected C2C12 cell lines were seeded on #1.5 glass coverslip (Cat. # 72230-01, Electron Microscopy Sciences) and labelled with cell non-permeable SNAP-Surface Alexa Fluor 488 substrate, incubated with 0, 0.2 or 20 nM human insulin for 30 min, fixed with 4% (w/v) paraformaldehyde (Sigma-Aldrich) for 20 minutes at room temperature, and washed 3 times with PBS. After permeabilized in 0.1% (v/v) Triton X-100 (Sigma-Aldrich) for 15 minutes, primary antibodies targeting Caveolin1 (1:400, Rabbit mAb, Cat. #3267, Cell Signaling, RRID: AB_2275453) and CLTC heavy chain (1:200, Goat pAb, Cat. #sc-6579, Santa Cruz, RRID:AB_2083170) were incubated overnight at 4°C. Similarly, for AXNA2 experiment, cells labeled with SNAP-Surface Alexa Fluor 488 substrate were incubated with insulin for 15 or 30 min, and labeled with Annexin A2 primary antibody (1:150, Rabbit mAb, Cat. # 8235S, Cell Signaling, RRID:AB_11129437). Coverslips were then washed in PBS, blocked in Dako Protein Block (Cat #X0909, Agilent), and stained by secondary antibodies (1:1000, Cy3 Donkey anti-Rabbit, Cat. #711-165-152, Jackson ImmunoResearch, RRID:AB_2307443; 1:500, Alexa Fluor 633 Donkey anti-Goat, Cat. # A21082, Thermo Fisher Scientific, RRID:AB_2535739) diluted in Dako Antibody Diluent (Cat. # S0809, Agilent) for 2 hours at room temperature. After washing in PBS, coverslips were mounted with Prolong Diamond Antifade Mountant (Cat. #P36961, Invitrogen).

### STED microscopy and image analysis

Stimulated emission depletion (STED) microscopy was performed with the 100×/1.4 Oil HC PL APO CS2 STED White objective of a Leica TCS SP8 3× STED microscope (Leica, Wetzlar, Germany) equipped with a white light laser, HyD detectors, and Leica Application Suite X (LAS X) software (LSI Imaging Core, Life Sciences Institute, University of British Columbia). Time-gated fluorescence detection was used to further improve lateral resolution. INSR-SNAP labelled by Alexa Fluor 488 was excited at 488 nm and depleted using the 592-nm depletion laser. CAV1 or AXNA2 labelled by Cy3 was excited at 555 nm and depleted using the 660 nm depletion laser. CLTC labelled by Alex Fluor 633 was excited at 633 nm and depleted using the 775-nm depletion laser. Sequential acquisition in the order of AF633/Cy3/AF488 was used to avoid cross-talk.

Huygens Professional software (Scientific Volume Imaging, Hilversum, the Netherlands) was used to deconvolve STED images and calculate the Object Pearson Coefficients for the colocalizations between INSR-SNAP, CAV1, CLTC, or ANXA2.

### TIRF microscopy and single particle tracking analysis

Total internal reflection fluorescence (TIRF) microscopy was performed on a Zeiss Axiovert 200M with a 100x Alpha-PlanFluar NA 1.45 oil objective (Zeiss, Germany) and a TIRF laser angle modifier (Zeiss). Cells were kept at 37°C in a temperature-controlled incubation chamber (Harvard Apparatus, Holliston, MA) and imaged in Ringer’s buffer supplemented with 20 mM glucose, as previously described(4). For insulin stimulations, cells were in the Ringer’s (20 mM glucose) buffer containing 0 (control) or 2 nM of insulin. Images were acquired with a CoolSNAP HQ2 CCD camera (Photometrics, Tucson, AZ). 405 nm and 561 nm solid-state diode laser system was used to excite TagBFP and mRFP tagged proteins.

The ImageJ distribution Fiji was used to process microscope images(83). The local background of TIRF images was subtracted using the rolling ball method (20-pixel radius) before single particle tracking.

Single particles (INSR, CAV1, or CLTC) were detected and tracked using Icy bioimaging analysis software version 1.9.8.1(84). Particles were detected by the UnDecimated Wavelet Transform Detector plugin that was set to detect bright spots over a dark background, exhibit 70% sensitivity to spots <3 pixels, detect particle size between 2 and 3000 pixels. Particles were tracked using the Multiple Hypothesis Tracking method built into the Spot Tracking plugin that was set to recognize particles that exhibited either diffusive or directed motion(85,86). Track radius (maximum displacement of the particle track) and lifetime of the particles were exported from the Motion Profile Processor of the Spot Tracking plugin. Diffusion coefficients were calculated in MATLAB as described previously but without corrections for positional errors and blurring(87). Colocalization of tracks was determined by Track Processor Interaction Analysis (distance threshold 2 pixels) in the Spot Tracking plugin. INSR tracks that colocalized with CAV1 or CLTC tracks for more than 4 continuous frames are determined to be colocalized tracks. The exported time and duration of colocalization were organized in Python and plotted in R studio. Diffusion coefficients, track radius, and the lifetime of tracks were plotted in R studio. Specifically, SuperPlots were used to plot the mean value of the cells and individual tracks of each cell(88). Code is available at GitHub deposit https://github.com/hcen/INSR_tracking.

### Skeletal muscle co-immunoprecipitation

Animal protocols were approved by the University of British Columbia Animal Care Committee in accordance with national guidelines. Mice received 1.5U/kg insulin (a dose commonly used in insulin tolerance test) or PBS through intraperitoneal injection. Skeletal muscle (gastrocnemius) was collected after 5 or 10 min of injection. INSR was immunoprecipitated by incubating 500 µg of protein lysate with 2 µg INSR antibodies (Rabbit pAb, Cat. #sc-711, Santa Cruz, RRID: AB_631835) at 4°C overnight. Subsequently, the solution was incubated with PureProteome™ Protein G Magnetic Beads (Cat. # LSKMAGG10, Millipore, Burlington, MA, USA) and washed according to the manufacturer’s instructions. INSR protein complexes were eluted from the beads by incubation in Blue Loading Buffer (50 mM DTT, Cell Signaling) at 95°C for 5 min and were analyzed by standard western blot procedures. INSR, CAV1, and CLTC were detected by the following Cell Signaling primary antibodies: INSRβ (1:1000, Mouse mAb, Cat. #3020S, RRID: AB_2249166), CLTC (1:1000, Rabbit mAb, Cat. #4796, RRID:AB_10828486), Caveolin1 (1:1000, Rabbit mAb, Cat. #3267, RRID: AB_2275453), and HRP-conjugated VeriBlot secondary antibodies (1:200, anti-mouse Cat. #ab131368, RRID:AB_2895114; anti-rabbit ab131366, RRID:AB_2892718; Abcam) that do not bind to IgG heavy and light chains to reduce background noise.

### Immunoprecipitation-Mass Spectrometry

C2C12 myoblasts were serum-starved for 6 hours and were given 0 or 2 nM insulin for 15 min before collecting their lysates in RIPA buffer (50 mM β-glycerol phosphate, 10 mM HEPES, 1% Triton X-100, 70 mM NaCl, 2 mM EGTA, 1 mM Na3VO4, and 1 mM NaF) supplemented with complete mini protease inhibitor cocktail (Roche, Laval, QC). INSRα antibody (NNC0276-3000, a gift from Novo Nordisk compound sharing program) that does not cross-react with IGF1R(34) was covalently conjugated to Dynabeads M-270 Epoxy beads (Cat. #14301, Invitrogen) to immunoprecipitate INSR according to the manufacturer’s protocol. INSR protein complexes were eluted from the beads by incubation in Blue Loading Buffer (50 mM DTT, Cell Signaling) at 95°C for 5 min.

Eluted co-immunoprecipitation samples were run into 10% SDS PAGE gel for clean-up procedure, fixed, stained, and destained in water. Each of the samples was excised, reduced, alkylated, and digested according to standard in-gel digest protocols(89). Extracted peptide samples were then cleaned up via STAGE-tip purification, as previously described(90).

Samples were reconstituted in 2% ACN, 0.5% formic acid, and the peptides were analyzed using a quadrupole – time of flight mass spectrometer (Impact II; Bruker Daltonics) on-line coupled to an EasyLC 1000 HPLC (ThermoFisher Scientific) using a Captive spray nanospray ionization source (Bruker Daltonics) including Aurora Series Gen2 (CSI) analytical column, (25cm x 75μm 1.6μm FSC C18, with Gen2 nanoZero and CSI fitting; Ion Opticks, Parkville, Victoria, Australia), and a μ-Precolumn, 300 μm ID x 5 mm, C18 PepMap, 5 μm, 100 A, Thermo Scientific, Waltham, Massachusetts, United States). Samples were reconstituted in (0.1% aqueous formic acid and 2 % acetonitrile in water) and 1ug of sample was loaded, injected in triplicates. A standard 90 min reversed-phase separation was employed, using water:acetonitrile gradients.

LC and MS acquisition parameters were standard for peptide sequencing. Acquired data were then searched by MaxQuant (V1.6.7.) against the Uniprot protein database for Mouse C2C12 cell line, and LFQ intensities extracted and normalized using the MaxLFQ algorithm, with 20ppm and 30ppm mass accuracies for precursor and product ion masses, respectively, and a 1% false discovery rate cut-off. Differential expression was determined by comparing the mean protein intensities of the triplicate injections between the groups. Significance (p<0.05) was calculated via t-test using default parameters in Perseus (v1.6.14.0).

### AlphaFold-Multimer and minD

Analysis of direct interactions between INSR and interactors was carried out using AlphaFold-Multimer and minD method(44,45). We downloaded all the protein sequences from UniProt (IDs: INSR - P15208, RPSA - P14206, HSPB1 - P14602, ANXA2 - P07356, HIST1H1B - P43276 and IGF1R - E9QNX9, CAV1 - P49817, CAV3 - P51637, AP2M1 - P84091, IRS1 - P35569, MTCO2 - P00405). We then followed the previously described protocol(45) to extract potential binding sites and do an in-silico evaluation of the interactions between binding sites. Briefly, we first paired INSR with other proteins and ran AlphaFold-Multimer on those pairs. We used full-length protein sequences to extract minD scores. minD is defined as the minimum expected distance of one residue of one protein chain from the other protein chain in the pairs. This metric is extracted from the Distogram head of AlphaFold-Multimer and has been shown to peak when a potential binding site is detected. After the binding sites on both proteins are identified, we then cut the region surrounding them. This results in a handful of fragments for each pair. We then pair fragments for a second round of AlphaFold-Multimer runs. Finally, ipTM scores are extracted to filter out non-binding pairs. A threshold of 0.42 is used to discriminate non-interacting and interacting fragment pairs(45). The code for this analysis is available at https://github.com/alirezaomidi/AFminD.

### Statistics

Data were presented as mean ± SEM in addition to the individual data points. A significance level of p < 0.05 was used throughout. Student’s t-tests were used for comparisons between two groups. Comparisons between more than two groups were performed using one-way ANOVA and multiple comparison p-values adjusted using the Tukey method. Comparisons between multiple groups and two categorical variables were performed using Two-way ANOVA followed by Tukey post hoc test. The statistical analysis above was performed using GraphPad Prism (Version 9.0.2). Two-sided Kolmogorov-Smirnov Tests were used for comparing cumulative probability, which was performed with ks.text() function in R.

## Data availability

Data are contained within the article or can be shared upon request.

## Acknowledgments

Spinning disk confocal and STED microscopy were performed in the LSI IMAGING facility of the Life Sciences Institute of the University of British Columbia using infrastructure funded by the Canadian Foundation of Innovation and BC Knowledge Development Fund as well as a Strategic Investment Fund (Faculty of Medicine, University of British Columbia). We thank Dr. Ivan Robert Nabi at UBC for guidance on imaging experiments and helpful discussions. Mass spectrometry used here was supported by the Canada Foundation for Innovation and Genome Canada (264PRO). We thank Yuanyuan Amanda Yang for guidance on Python and R coding, Yi Han Xia for assistance in R coding, and Jiashuo Aaron Zhang for assistance in data analysis. We thank Michael G. Atser for assistance in western blots. INSR antibody NNC0276-3000 was provided by Novo Nordisk Compound Sharing. The schematics in Figure 7 were created with BioRender.

## Notes

Conflict of interest: The authors declare that they have no conflicts of interest with the contents of this article.

https://doi.org/10.5281/zenodo.18272839

### Competing Interest Statement

The authors have declared no competing interest.

### Summary of Updates

Minor edits on the title and main text. Added RRID in the methods.

## References

1. Knutson VP. Cellular trafficking and processing of the insulin receptor. FASEB J. 1991;5(8):2130–2138.

2. Kahn CR, Crettaz M. Insulin receptors and the molecular mechanism of insulin action. Diabetes Metab Rev. 1985;1(1-2):5–32.

3. Bevan AP, Burgess JW, Drake PG, Shaver A, Bergeron JJ, Posner BI. Selective activation of the rat hepatic endosomal insulin receptor kinase. Role for the endosome in insulin signaling. The Journal of biological chemistry. 1995;270(18):10784–10791.

4. Boothe T, Lim GE, Cen H, Skovsø S, Piske M, Li SN, Nabi IR, Gilon P, Johnson JD. Inter-domain tagging implicates caveolin-1 in insulin receptor trafficking and Erk signaling bias in pancreatic beta-cells. Mol Metab. 2016;5(5):366–378.

5. Trischitta V, Gullo D, Squatrito S, Pezzino V, Goldfine ID, Vigneri R. Insulin Internalization into Monocytes Is Decreased in Patients with Type II Diabetes Mellitus. 1986;62:522–528.

6. Trischitta V, Brunetti A, Chiavetta A, Benzi L, Papa V, Vigneri R. Defects in insulin–receptor internalization and processing in monocytes of obese subjects and obese NIDDM patients. 1989;38:1579–1584.

7. Jochen AL, Berhanu P, Olefsky JM. Insulin internalization and degradation in adipocytes from normal and type II diabetic subjects. J Clin Endocrinol Metab. 1986;62(2):268–274.

8. Rask-Madsen C, Kahn CR. Tissue-specific insulin signaling, metabolic syndrome, and cardiovascular disease. Arterioscler Thromb Vasc Biol. 2012;32(9):2052–2059.

9. Guo S. Insulin signaling, resistance, and the metabolic syndrome: insights from mouse models into disease mechanisms. J Endocrinol. 2014;220(2):T1–T23.

10. Jochen A, Hays J, Lee M. Kinetics of insulin internalization and processing in adipocytes: effects of insulin concentration. J Cell Physiol. 1989;141(3):527–534.

11. McClain DA, Olefsky JM. Evidence for two independent pathways of insulin-receptor internalization in hepatocytes and hepatoma cells. Diabetes. 1988;37(6):806–815.

12. Ceresa BP, Kao AW, Santeler SR, Pessin JE. Inhibition of clathrin-mediated endocytosis selectively attenuates specific insulin receptor signal transduction pathways. Mol Cell Biol. 1998;18(7):3862–3870.

13. Carpentier JL, Van Obberghen E, Gorden P, Orci L. Surface redistribution of 125I-insulin in cultured human lymphocytes. J Cell Biol. 1981;91(1):17–25.

14. Paccaud JP, Siddle K, Carpentier JL. Internalization of the human insulin receptor. The insulin-independent pathway. The Journal of biological chemistry. 1992;267(18):13101–13106.

15. Jose M, Biosca JA, Trujillo R, Itarte E. Characterization of the hepatic insulin receptor undergoing internalization through clathrin-coated vesicles and endosomes. FEBS Lett. 1993;334(3):286–288.

16. Nystrom FH, Chen H, Cong LN, Li Y, Quon MJ. Caveolin-1 interacts with the insulin receptor and can differentially modulate insulin signaling in transfected Cos-7 cells and rat adipose cells. Mol Endocrinol. 1999;13(12):2013–2024.

17. Fagerholm S, Ortegren U, Karlsson M, Ruishalme I, Str√•lfors P. Rapid insulin-dependent endocytosis of the insulin receptor by caveolae in primary adipocytes. PLoS ONE. 2009;4(6):e5985.

18. Knowler WC, Fowler SE, Hamman RF, Christophi CA, Hoffman HJ, Brenneman AT, Brown-Friday JO, Goldberg R, Venditti E, Nathan DM, Group DPPR. 10-year follow-up of diabetes incidence and weight loss in the Diabetes Prevention Program Outcomes Study. Lancet. 2009;374(9702):1677–1686.

19. Moore MC, Cherrington AD, Wasserman DH. Regulation of hepatic and peripheral glucose disposal. Best Pract Res Clin Endocrinol Metab. 2003;17(3):343–364.

20. Tseng LT, Lin CL, Tzen KY, Chang SC, Chang MF. LMBD1 protein serves as a specific adaptor for insulin receptor internalization. The Journal of biological chemistry. 2013;288(45):32424–32432.

21. Gautier A, Juillerat A, Heinis C, Corrêa IR, Kindermann M, Beaufils F, Johnsson K. An engineered protein tag for multiprotein labeling in living cells. Chem Biol. 2008;15(2):128–136.

22. Melmed S, Polonsky KS, Larsen PR, Kronenberg HM. Williams Textbook of Endocrinology. 13th ed: Elsevier Saunders.

23. Timmons JA, Atherton PJ, Larsson O, Sood S, Blokhin IO, Brogan RJ, Volmar CH, Josse AR, Slentz C, Wahlestedt C, Phillips SM, Phillips BE, Gallagher IJ, Kraus WE. A coding and non-coding transcriptomic perspective on the genomics of human metabolic disease. Nucleic Acids Res. 2018;46(15):7772–7792.

24. McAuley KA, Williams SM, Mann JI, Walker RJ, Lewis-Barned NJ, Temple LA, Duncan AW. Diagnosing insulin resistance in the general population. Diabetes Care. 2001;24(3):460–464.

25. Hengist A, Edinburgh RM, Davies RG, Walhin JP, Buniam J, James LJ, Rogers PJ, Gonzalez JT, Betts JA. Physiological responses to maximal eating in men. Br J Nutr. 2020;124(4):407–417.

26. Polonsky KS, Given BD, Hirsch LJ, Tillil H, Shapiro ET, Beebe C, Frank BH, Galloway JA, Van Cauter E. Abnormal patterns of insulin secretion in non-insulin-dependent diabetes mellitus. N Engl J Med. 1988;318(19):1231–1239.

27. Hamley S, Kloosterman D, Duthie T, Dalla Man C, Visentin R, Mason SA, Ang T, Selathurai A, Kaur G, Morales-Scholz MG, Howlett KF, Kowalski GM, Shaw CS, Bruce CR. Mechanisms of hyperinsulinaemia in apparently healthy non-obese young adults: role of insulin secretion, clearance and action and associations with plasma amino acids. Diabetologia. 2019;62(12):2310–2324.

28. Le Marchand-Brustel Y, Jeanrenaud B, Freychet P. Insulin binding and effects in isolated soleus muscle of lean and obese mice. Am J Physiol. 1978;234(4):E348–358.

29. Cen HH, Hussein B, Botezelli JD, Wang S, Zhang JA, Noursadeghi N, Jessen N, Rodrigues B, Timmons JA, Johnson JD. Human and mouse muscle transcriptomic analyses identify insulin receptor mRNA downregulation in hyperinsulinemia-associated insulin resistance. FASEB J. 2022;36(1):e22088.

30. Cen HH, Johnson JD. Insulin receptor trafficking and interactions in muscle cells - Supplemental Materials. Zenodo. 2026; 10.5281/zenodo.18272839

31. Jans DA. Lateral Mobility of Polypeptide Hormone Receptors and GTP-Binding Proteins. The Mobile Receptor Hypothesis: The Role of Membrane Receptor Lateral Movement in Signal Transduction. Boston, MA: Springer US; 1997:83–115.

32. Kabayama K, Sato T, Saito K, Loberto N, Prinetti A, Sonnino S, Kinjo M, Igarashi Y, Inokuchi J. Dissociation of the insulin receptor and caveolin-1 complex by ganglioside GM3 in the state of insulin resistance. Proc Natl Acad Sci U S A. 2007;104(34):13678–13683.

33. Ikonen E, Vainio S. Lipid microdomains and insulin resistance: is there a connection? Sci STKE. 2005;2005(268):pe3.

34. Ørstrup LH, Slaaby R, Rasch MG, Rasmussen N, Lund S, Brandt J, Schluckebier G, Wang Z, Lützen A, Pedersen T, Hvid H, Hansen BF, Blume N. Cross-species reactive monoclonal antibodies against the extracellular domains of the insulin receptor and IGF1 receptor. J Immunol Methods. 2019;465:20–26.

35. Benyoucef S, Surinya KH, Hadaschik D, Siddle K. Characterization of insulin/IGF hybrid receptors: contributions of the insulin receptor L2 and Fn1 domains and the alternatively spliced exon 11 sequence to ligand binding and receptor activation. Biochem J. 2007;403(3):603–613.

36. Moxham CP, Duronio V, Jacobs S. Insulin-like growth factor I receptor beta-subunit heterogeneity. Evidence for hybrid tetramers composed of insulin-like growth factor I and insulin receptor heterodimers. The Journal of biological chemistry. 1989;264(22):13238–13244.

37. Treadway JL, Morrison BD, Goldfine ID, Pessin JE. Assembly of insulin/insulin-like growth factor-1 hybrid receptors in vitro. J Biol Chem. 1989;264(36):21450–21453.

38. Shu L, Lee L, Chang Y, Holzman LB, Edwards CA, Shelden E, Shayman JA. Caveolar structure and protein sorting are maintained in NIH 3T3 cells independent of glycosphingolipid depletion. Arch Biochem Biophys. 2000;373(1):83–90.

39. DiGiacomo V, Meruelo D. Looking into laminin receptor: critical discussion regarding the non-integrin 37/67-kDa laminin receptor/RPSA protein. Biol Rev Camb Philos Soc. 2016;91(2):288–310.

40. Zoubeidi A, Zardan A, Wiedmann RM, Locke J, Beraldi E, Fazli L, Gleave ME. Hsp27 promotes insulin-like growth factor-I survival signaling in prostate cancer via p90Rsk-dependent phosphorylation and inactivation of BAD. Cancer Res. 2010;70(6):2307–2317.

41. Muranova LK, Shatov VM, Bukach OV, Gusev NB. Cardio-Vascular Heat Shock Protein (cvHsp, HspB7), an Unusual Representative of Small Heat Shock Protein Family. Biochemistry (Mosc). 2021;86(Suppl 1):S1–S11.

42. Yuan H, Wang T, Niu Y, Liu X, Fu L. AMP-activated protein kinase-mediated expression of heat shock protein beta 1 enhanced insulin sensitivity in the skeletal muscle. FEBS Lett. 2017;591(1):97–108.

43. Jumper J, Evans R, Pritzel A, Green T, Figurnov M, Ronneberger O, Tunyasuvunakool K, Bates R, Žídek A, Potapenko A, Bridgland A, Meyer C, Kohl SAA, Ballard AJ, Cowie A, Romera-Paredes B, Nikolov S, Jain R, Adler J, Back T, Petersen S, Reiman D, Clancy E, Zielinski M, Steinegger M, Pacholska M, Berghammer T, Bodenstein S, Silver D, Vinyals O, Senior AW, Kavukcuoglu K, Kohli P, Hassabis D. Highly accurate protein structure prediction with AlphaFold. Nature. 2021;596(7873):583–589.

44. Zhu W, Shenoy A, Kundrotas P, Elofsson A. Evaluation of AlphaFold-Multimer prediction on multi-chain protein complexes. Bioinformatics. 2023;39(7).

45. Omidi A, Møller MH, Malhis N, Bui JM, Gsponer J. AlphaFold-Multimer accurately captures interactions and dynamics of intrinsically disordered protein regions. Proc Natl Acad Sci U S A. 2024;121(44):e2406407121.

46. Chidlow JH, Sessa WC. Caveolae, caveolins, and cavins: complex control of cellular signalling and inflammation. Cardiovasc Res. 2010;86(2):219–225.

47. Capozza F, Cohen AW, Cheung MW, Sotgia F, Schubert W, Battista M, Lee H, Frank PG, Lisanti MP. Muscle-specific interaction of caveolin isoforms: differential complex formation between caveolins in fibroblastic vs. muscle cells. Am J Physiol Cell Physiol. 2005;288(3):C677–691.

48. Schöneich C, Dremina E, Galeva N, Sharov V. Apoptosis in differentiating C2C12 muscle cells selectively targets Bcl-2-deficient myotubes. Apoptosis. 2014;19(1):42–57.

49. Biener Y, Feinstein R, Mayak M, Kaburagi Y, Kadowaki T, Zick Y. Annexin II is a novel player in insulin signal transduction. Possible association between annexin II phosphorylation and insulin receptor internalization. The Journal of biological chemistry. 1996;271(46):29489–29496.

50. Rescher U, Ludwig C, Konietzko V, Kharitonenkov A, Gerke V. Tyrosine phosphorylation of annexin A2 regulates Rho-mediated actin rearrangement and cell adhesion. J Cell Sci. 2008;121(Pt 13):2177–2185.

51. Carpentier JL. Insulin-induced and constitutive internalization of the insulin receptor. Horm Res. 1992;38(1-2):13–18.

52. Chang M, Kwon M, Kim S, Yunn NO, Kim D, Ryu SH, Lee JB. Aptamer-based single-molecule imaging of insulin receptors in living cells. J Biomed Opt. 2014;19(5):051204.

53. Hubert M, Larsson E, Lundmark R. Keeping in touch with the membrane; protein-and lipid-mediated confinement of caveolae to the cell surface. Biochem Soc Trans. 2020;48(1):155–163.

54. Martins AS, Ordóñez JL, Amaral AT, Prins F, Floris G, Debiec-Rychter M, Hogendoorn PC, de Alava E. IGF1R signaling in Ewing sarcoma is shaped by clathrin-/caveolin-dependent endocytosis. PLoS One. 2011;6(5):e19846.

55. Pandini G, Frasca F, Mineo R, Sciacca L, Vigneri R, Belfiore A. Insulin/insulin-like growth factor I hybrid receptors have different biological characteristics depending on the insulin receptor isoform involved. The Journal of biological chemistry. 2002;277(42):39684–39695.

56. Tomas A, Futter CE, Eden ER. EGF receptor trafficking: consequences for signaling and cancer. Trends Cell Biol. 2014;24(1):26–34.

57. Viola CM, Frittmann O, Jenkins HT, Shafi T, De Meyts P, Brzozowski AM. Structural conservation of insulin/IGF signalling axis at the insulin receptors level in Drosophila and humans. Nature Communications. 2023;14(1):6271.

58. Kammanadiminti SJ, Chadee K. Suppression of NF-kappaB activation by Entamoeba histolytica in intestinal epithelial cells is mediated by heat shock protein 27. The Journal of biological chemistry. 2006;281(36):26112–26120.

59. Ku HC, Chang HH, Liu HC, Hsiao CH, Lee MJ, Hu YJ, Hung PF, Liu CW, Kao YH. Green tea (-)-epigallocatechin gallate inhibits insulin stimulation of 3T3-L1 preadipocyte mitogenesis via the 67-kDa laminin receptor pathway. Am J Physiol Cell Physiol. 2009;297(1):C121–132.

60. Zhao WQ, Chen GH, Chen H, Pascale A, Ravindranath L, Quon MJ, Alkon DL. Secretion of Annexin II via activation of insulin receptor and insulin-like growth factor receptor. The Journal of biological chemistry. 2003;278(6):4205–4215.

61. Caron D, Boutchueng-Djidjou M, Tanguay RM, Faure RL. Annexin A2 is SUMOylated on its N-terminal domain: regulation by insulin. FEBS Lett. 2015;589(9):985–991.

62. Urbanska A, Sadowski L, Kalaidzidis Y, Miaczynska M. Biochemical characterization of APPL endosomes: the role of annexin A2 in APPL membrane recruitment. Traffic. 2011;12(9):1227–1241.

63. Ryu J, Galan AK, Xin X, Dong F, Abdul-Ghani MA, Zhou L, Wang C, Li C, Holmes BM, Sloane LB, Austad SN, Guo S, Musi N, DeFronzo RA, Deng C, White MF, Liu F, Dong LQ. APPL1 potentiates insulin sensitivity by facilitating the binding of IRS1/2 to the insulin receptor. Cell Rep. 2014;7(4):1227–1238.

64. Lemmon MA, Schlessinger J. Cell signaling by receptor tyrosine kinases. Cell. 2010;141(7):1117–1134.

65. de Graauw M, Cao L, Winkel L, van Miltenburg MH, le Dévédec SE, Klop M, Yan K, Pont C, Rogkoti VM, Tijsma A, Chaudhuri A, Lalai R, Price L, Verbeek F, van de Water B. Annexin A2 depletion delays EGFR endocytic trafficking via cofilin activation and enhances EGFR signaling and metastasis formation. Oncogene. 2014;33(20):2610–2619.

66. Behrends M, Engmann O. Linker histone H1.5 is an underestimated factor in differentiation and carcinogenesis. Environ Epigenet. 2020;6(1):dvaa013.

67. Brown M, Dainty S, Strudwick N, Mihai AD, Watson JN, Dendooven R, Paton AW, Paton JC, Schröder M. Endoplasmic reticulum stress causes insulin resistance by inhibiting delivery of newly synthesized insulin receptors to the cell surface. Mol Biol Cell. 2020;31(23):2597–2629.

68. Vicidomini G, Bianchini P, Diaspro A. STED super-resolved microscopy. Nat Methods. 2018;15(3):173–182.

69. Fish KN. Total internal reflection fluorescence (TIRF) microscopy. Curr Protoc Cytom. 2009;Chapter 12:Unit12.18.

70. Le PU, Guay G, Altschuler Y, Nabi IR. Caveolin-1 is a negative regulator of caveolae-mediated endocytosis to the endoplasmic reticulum. Journal of Biological Chemistry. 2002;277(5):3371–3379.

71. Humphries AC, Way M. The non-canonical roles of clathrin and actin in pathogen internalization, egress and spread. Nat Rev Microbiol. 2013;11(8):551–560.

72. Rodal SK, Skretting G, Garred O, Vilhardt F, van Deurs B, Sandvig K. Extraction of cholesterol with methyl-beta-cyclodextrin perturbs formation of clathrin-coated endocytic vesicles. Mol Biol Cell. 1999;10(4):961–974.

73. Parpal S, Karlsson M, Thorn H, Strålfors P. Cholesterol depletion disrupts caveolae and insulin receptor signaling for metabolic control via insulin receptor substrate-1, but not for mitogen-activated protein kinase control. The Journal of biological chemistry. 2001;276(13):9670–9678.

74. Cohen AW, Combs TP, Scherer PE, Lisanti MP. Role of caveolin and caveolae in insulin signaling and diabetes. Am J Physiol Endocrinol Metab. 2003;285(6):E1151–1160.

75. Hussain KM, Leong KL, Ng MM, Chu JJ. The essential role of clathrin-mediated endocytosis in the infectious entry of human enterovirus 71. The Journal of biological chemistry. 2011;286(1):309–321.

76. Wu J, Park SH, Choi E. The insulin receptor endocytosis. Prog Mol Biol Transl Sci. 2023;194:79–107.

77. Chen Y, Huang L, Qi X, Chen C. Insulin Receptor Trafficking: Consequences for Insulin Sensitivity and Diabetes. Int J Mol Sci. 2019;20(20).

78. Choi E, Zhang X, Xing C, Yu H. Mitotic Checkpoint Regulators Control Insulin Signaling and Metabolic Homeostasis. Cell. 2016;166(3):567–581.

79. Park J, Hall C, Hubbard B, LaMoia T, Gaspar R, Nasiri A, Li F, Zhang H, Kim J, Haeusler RA, Accili D, Shulman GI, Yu H, Choi E. MAD2-Dependent Insulin Receptor Endocytosis Regulates Metabolic Homeostasis. Diabetes. 2023;72(12):1781–1794.

80. Choi E, Kikuchi S, Gao H, Brodzik K, Nassour I, Yopp A, Singal AG, Zhu H, Yu H. Mitotic regulators and the SHP2-MAPK pathway promote IR endocytosis and feedback regulation of insulin signaling. Nat Commun. 2019;10(1):1473.

81. Ansarullah, Jain C, Far FF, Homberg S, Wißmiller K, von Hahn FG, Raducanu A, Schirge S, Sterr M, Bilekova S, Siehler J, Wiener J, Oppenländer L, Morshedi A, Bastidas-Ponce A, Collden G, Irmler M, Beckers J, Feuchtinger A, Grzybek M, Ahlbrecht C, Feederle R, Plettenburg O, Müller TD, Meier M, Tschöp MH, Coskun Ü, Lickert H. Inceptor counteracts insulin signalling in β-cells to control glycaemia. Nature. 2021;590(7845):326–331.

82. Siehler J, Bilekova S, Chapouton P, Dema A, Albanese P, Tamara S, Jain C, Sterr M, Enos SJ, Chen C, Malhotra C, Villalba A, Schomann L, Bhattacharya S, Feng J, Akgün Canan M, Ribaudo F, Ansarullah, Burtscher I, Ahlbrecht C, Plettenburg O, Kurth T, Scharfmann R, Speier S, Scheltema RA, Lickert H. Inceptor binds to and directs insulin towards lysosomal degradation in βLcells. Nat Metab. 2024;6(12):2374–2390.

83. Schindelin J, Arganda-Carreras I, Frise E, Kaynig V, Longair M, Pietzsch T, Preibisch S, Rueden C, Saalfeld S, Schmid B, Tinevez JY, White DJ, Hartenstein V, Eliceiri K, Tomancak P, Cardona A. Fiji: an open-source platform for biological-image analysis. Nat Methods. 2012;9(7):676–682.

84. de Chaumont F, Dallongeville S, Chenouard N, Hervé N, Pop S, Provoost T, Meas-Yedid V, Pankajakshan P, Lecomte T, Le Montagner Y, Lagache T, Dufour A, Olivo-Marin JC. Icy: an open bioimage informatics platform for extended reproducible research. Nat Methods. 2012;9(7):690–696.

85. Chenouard N, Bloch I, Olivo-Marin JC. Multiple hypothesis tracking for cluttered biological image sequences. IEEE Trans Pattern Anal Mach Intell. 2013;35(11):2736–3750.

86. Chenouard N, Smal I, de Chaumont F, Maška M, Sbalzarini IF, Gong Y, Cardinale J, Carthel C, Coraluppi S, Winter M, Cohen AR, Godinez WJ, Rohr K, Kalaidzidis Y, Liang L, Duncan J, Shen H, Xu Y, Magnusson KE, Jaldén J, Blau HM, Paul-Gilloteaux P, Roudot P, Kervrann C, Waharte F, Tinevez JY, Shorte SL, Willemse J, Celler K, van Wezel GP, Dan HW, Tsai YS, Ortiz de Solórzano C, Olivo-Marin JC, Meijering E. Objective comparison of particle tracking methods. Nat Methods. 2014;11(3):281–289.

87. Abraham L, Lu HY, Falcão RC, Scurll J, Jou T, Irwin B, Tafteh R, Gold MR, Coombs D. Limitations of Qdot labelling compared to directly-conjugated probes for single particle tracking of B cell receptor mobility. Sci Rep. 2017;7(1):11379.

88. Lord SJ, Velle KB, Mullins RD, Fritz-Laylin LK. SuperPlots: Communicating reproducibility and variability in cell biology. J Cell Biol. 2020;219(6).

89. Shevchenko A, Wilm M, Vorm O, Mann M. Mass spectrometric sequencing of proteins silver-stained polyacrylamide gels. Anal Chem. 1996;68(5):850–858.

90. Rappsilber J, Mann M, Ishihama Y. Protocol for micro-purification, enrichment, pre-fractionation and storage of peptides for proteomics using StageTips. Nat Protoc. 2007;2(8):1896–1906.

